# The Hox code abolishes anatomical variation in an insect gonad

**DOI:** 10.64898/2026.10.09.752918

**Authors:** Jonchee A. Kao, Tarun Kumar, Daniella A. Haber, Cassandra G. Extavour

## Abstract

Phenotypic traits can vary based on genes or environment. Nevertheless, some quantitative traits remain invariant. Understanding the developmental mechanisms that generate such invariance can reveal how phenotypic robustness evolves. Insect reproductive capacity is largely determined by the number of gamete-producing subunits within the gonad, providing a tractable system to investigate these mechanisms as they operate on a quantitative trait that impacts fitness. In most insects, gonad subunit number varies within and between species. Here, we investigate gonad development in the milkweed bug *Oncopeltus fasciatus*, wherein subunit number is invariant. We show that subunit number is established during embryogenesis, when some abdominal segments, but not others, specify somatic gonad precursors. The position and number of these precursors depend on segment identity, and experimentally modified segment number or Hox-mediated segment identity changes the number of gonad subunits. Thus, *O. fasciatus* establishes invariant subunit number by coupling gonad development to segmental patterning machinery. We propose that co-option of the Hox code provides a developmental mechanism for generating robustness in gonad subunit number and may thereby contribute to the evolution of reproductive capacity in other insects.

## Introduction

Developmental processes can produce highly stereotypic and reproducible anatomical structures despite genetic, environmental, and stochastic variation ^1^. Standing variation is often considered a prerequisite for evolutionary change ^2,3^. Likewise, evolvability can correlate positively with adaptability ^4^, leading to the prediction that some level of trait variation should be favoured by natural selection ^5^. Nevertheless, some traits show remarkably little variation between or even within lineages ^6,7^, even traits that, if changed, might be predicted to increase organismal fitness ^8^.

Reproductive organ formation depends on multiple morphogenetic processes, including cell specification ^9–12^, migration ^13–15^, proliferation ^16,17^, and differentiation ^9^, each of which can vary within and between species ^18–27^. Understanding how developmental mechanisms buffer variation and generate consistent phenotypes is central to understanding such phenotypic robustness ^1^. Here we address this problem using a quantitative trait that can directly impact reproductive capacity and therefore fitness: insect ovariole and testiole number.

All insects have paired gonads composed of functional subunits called ovarioles in females and testioles, testicular follicles, or sperm tubes in males ^28–30^. In this work, we collectively call these subunits “gonadioles”. Between species, the number of gonadioles varies from one to many hundreds per gonad ^8^ (Figure 1A). Within species, gonadiole number usually also exhibits species-specific variation, on the order of 10% of the mean ^8,28,31^. However, some species show invariant gonadiole numbers ^8^ (Figure 1A), providing an example of phenotypic robustness. Gonadiole number therefore provides a tractable system for investigating how developmental mechanisms can either promote phenotypic variation, or constrain it, yielding robustness.

**Figure 1.**
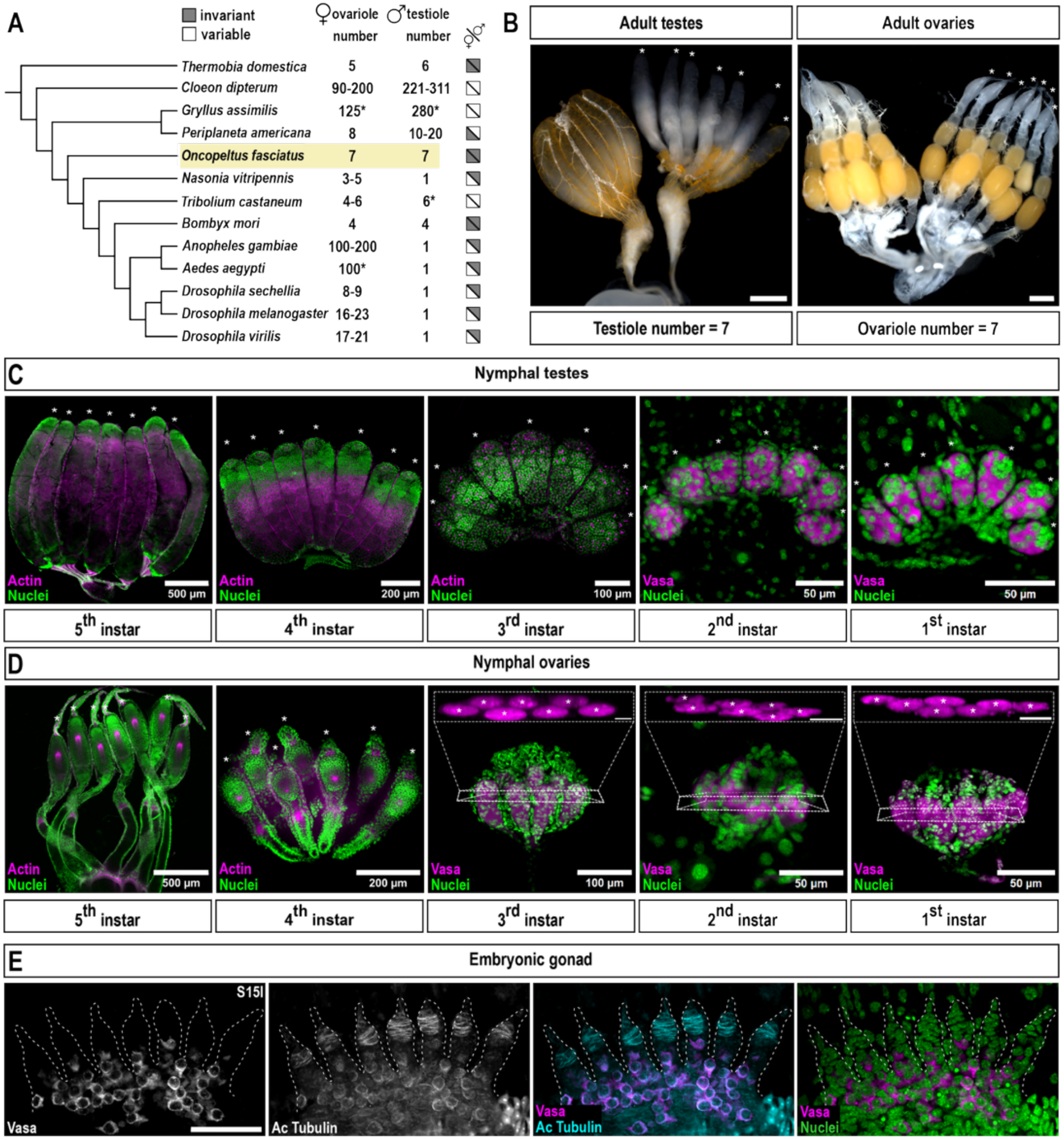
*O. fasciatus* gonadiole number is established in the embryo. **(A)** Cladogram illustrating the phylogenetic relationships amongst previously studied insect species. Ovariole number per ovary and testiole number per testis for each species (Table S1) are shown as a single value, a reported range (minimum – maximum), or an approximate mean (indicated by *). **(B)** Paired gonads of the adult hemipteran bug *O. fasciatus*. Gonads on the left are shown intact, whereas those on the right are dissected to reveal individual gonadioles. Asterisks denote each individual gonadiole, totalling seven ovarioles per ovary in females (n=20) and seven testioles per testis in males (n=20). Scale bars = 500 µm. **(C-D)** Gonad development across five nymphal stages. Individual testes (C) or ovaries (D) from each nymphal stage are shown, with asterisks denoting individual gonadioles. In later stage gonads, tissue architecture is visualised using phalloidin staining which labels filamentous actin (magenta). In earlier stage gonads, immunostaining against Ofas-Vas (magenta) is used to identify germ cells. Scale bar lengths are 500, 200, 100, 50 and 50 µm for 5^th^, 4^th^, 3^rd^, 2^nd^ and 1^st^ instar panels respectively. Insets above 3^rd^, 2^nd^ and 1^st^ instar ovary panels show transverse sections of the ovary where each ovariole is clearly resolved (inset scale bars = 20 µm). Nuclei are stained with Hoechst 33342 (green). **(E)** Embryonic gonad in a late-stage 15 embryo. Seven embryonic gonadiole primordia are outlined by dotted lines. Ofas-Vasa (magenta) labels the germ cells, while acetylated Tubulin (Ac Tubulin, cyan) highlights somatic gonadal structures and is detectable at low levels in the germ cells. Nuclei are stained with Hoechst 33342 (green). Scale bars = 50 µm.

Most developmental studies of gonadiole number have focused on ovariole number in *Drosophila* (Diptera) species of fruit flies. Amongst these species, mean ovariole number, though genetically regulated and therefore heritable ^18,32–39^, also shows within-species variation, including species-specific phenotypic plasticity with temperature ^40–42^, altitudinal and latitudinal clines^42–48^ and nutritional conditions ^31,41,49^.

The developmental regulation of ovariole number variation in *Drosophila melanogaster* begins with establishment of the embryonic gonad, which consists of both germ cells and somatic gonadal precursors (SGPs) ^9^. Primordial germ cells form at the posterior of the early embryo ^50^ and subsequently migrate through the embryo ^13^ to associate with SGPs derived principally from the mesoderm of parasegments 10–12 ^9,51^. These cells coalesce to form the embryonic gonad ^9,52,53^. During late larval development, a subset of somatic ovarian cells form multicellular structures called terminal filaments at the anterior of the ovary ^54,55^. Each terminal filament leads to the establishment of one mature ovariole, making larval terminal filament number a determinant of adult ovariole number ^55,56^.

Variation in adult ovariole number in *Drosophila* can be caused by developmental variation in almost any stage of ovarian development, including the specification and proliferation of somatic gonadal precursors ^19^, the allocation of somatic cells to the terminal-filament lineage ^18,19,41^, and the number of cells incorporated into each terminal filament ^38,41^. Differences in these processes contribute to ovariole-number variation within and between *Drosophila* lineages ^18,19,31,38,41^.

Much less is known about the developmental regulation of the invariant ovariole number seen in other insects. Classical embryological studies suggest that some ametabolous insects establish ovariole primordia in defined abdominal segments, such that the number of contributing segments corresponds to the adult ovariole number ^57,58^. Similarly, in the European fire bug *Pyrrhocoris apterus* (Hemiptera), eight pairs of gonadiole primordia reportedly form in abdominal segments A1–A8 ^59^. The A1 primordia subsequently degenerate, leaving seven gonadioles on each side ^59^. These observations suggest a relationship between segmental organization and gonadiole number, but the mechanisms that might specify the identity and number of gonad-forming segments remain unknown.

The milkweed bug *Oncopeltus fasciatus* provides an opportunity to investigate these mechanisms genetically. Adult females possess seven ovarioles per ovary, and adult males possess seven testioles per testis ^60,61^, with no variation in gonadiole number reported in the literature ^60,61^, nor observed in our laboratory over hundreds of adult dissections. The gonadioles in both sexes are reported to be established by hatching, suggesting that their number is determined during embryogenesis ^60,61^.

Here, we examine germline and somatic gonad development in *O. fasciatus*. We provide evidence that instead of using the *D. melanogaster* mechanism of variable postembryonic cellular organisation within the gonad, *O. fasciatus* co-opts embryonic segment identity programs to produce an invariant number of gonadioles per gonad. We propose that this developmental coupling of body plan and gonadal organization provides robustness to this reproductive trait, and may represent a more general mechanism for establishing invariance in organ system anatomies.

## Results

### Gonadiole number in Oncopeltus fasciatus is established during embryogenesis

Adult *Oncopeltus fasciatus* have seven ovarioles per ovary and seven testioles per testis (Figure 1A–B), with no intra-species variation in gonadiole number ^60,61^ (Tables S1, S4). To determine when adult gonadiole number is established during development, we examined gonads at progressively earlier developmental stages, identifying nymphal instars based on characteristic wing morphology ^62^. We observed seven testioles (Figure 1C) and seven ovarioles (Figure 1D) in all five nymphal instars, suggesting that gonadiole number is established during embryogenesis.

We next examined embryonic gonad development using previously defined developmental stages ^63^, further adapted for this study (Figure S1A; Table S2). By late stage 15 (approximately 60% of the way through embryogenesis, just over two days before hatching), seven distinct gonadiole primordia associated with primordial germ cells (PGCs) were evident (Figure 1E), confirming that the seven gonadioles are established during embryogenesis. We discovered that acetylated Tubulin, a post-translationally modified form of Tubulin associated with stable, long-lived microtubules ^64,65^, was enriched at the distal tips of each of the seven embryonic gonadiole primordia (Figure 1E). Using acetylated Tubulin as a marker of gonadioles, we then examined earlier embryonic stages. The earliest stage at which we detected acetylated Tubulin associated with these structures was stage 12, at which point individual gonadiole primordia were already distinguishable (Figure S1B, white arrowheads).

### Segment-specific mesodermal expression domains of O. fasciatus traffic jam and tinman orthologs form the somatic gonad precursors

To identify potential molecular markers for the gonadiole primordia at earlier stages, we examined the embryonic expression of the *O. fasciatus* homologs of genes expressed in or near the embryonic somatic gonad in *D. melanogaster* (Figure 2, Figure S1C–D, Table S3; also see Methods). In *D. melanogaster*, *tinman* (*tin*) expression is required for the specification of the somatic gonad precursors (SGPs) from the abdominal mesoderm of parasegments 10, 11 and 12 ^53,66^, and is also expressed in the dorsal vessel spanning segments T2–A6 ^67–69^. All embryonic SGPs in *D. melanogaster* express Traffic jam (Tj) protein ^70^, and Tj-positive somatic cells are intermingled with the germ cells throughout larval development ^70,71^. We found that in *O. fasciatus* late stage 15 (S15l) embryos, each putative gonadiole primordium appeared to consist of mesodermal regions with distinct gene expression profiles: a lateral domain expressing *Ofas*-*tin* and a medial domain expressing *Ofa*s-*tj* (Figure 2M–M”). To characterize the emergence and morphogenesis of these domains, we examined *Ofas*-*tin* and *Ofas*-*tj* expression throughout embryogenesis (Figure 2).

**Figure 2.**
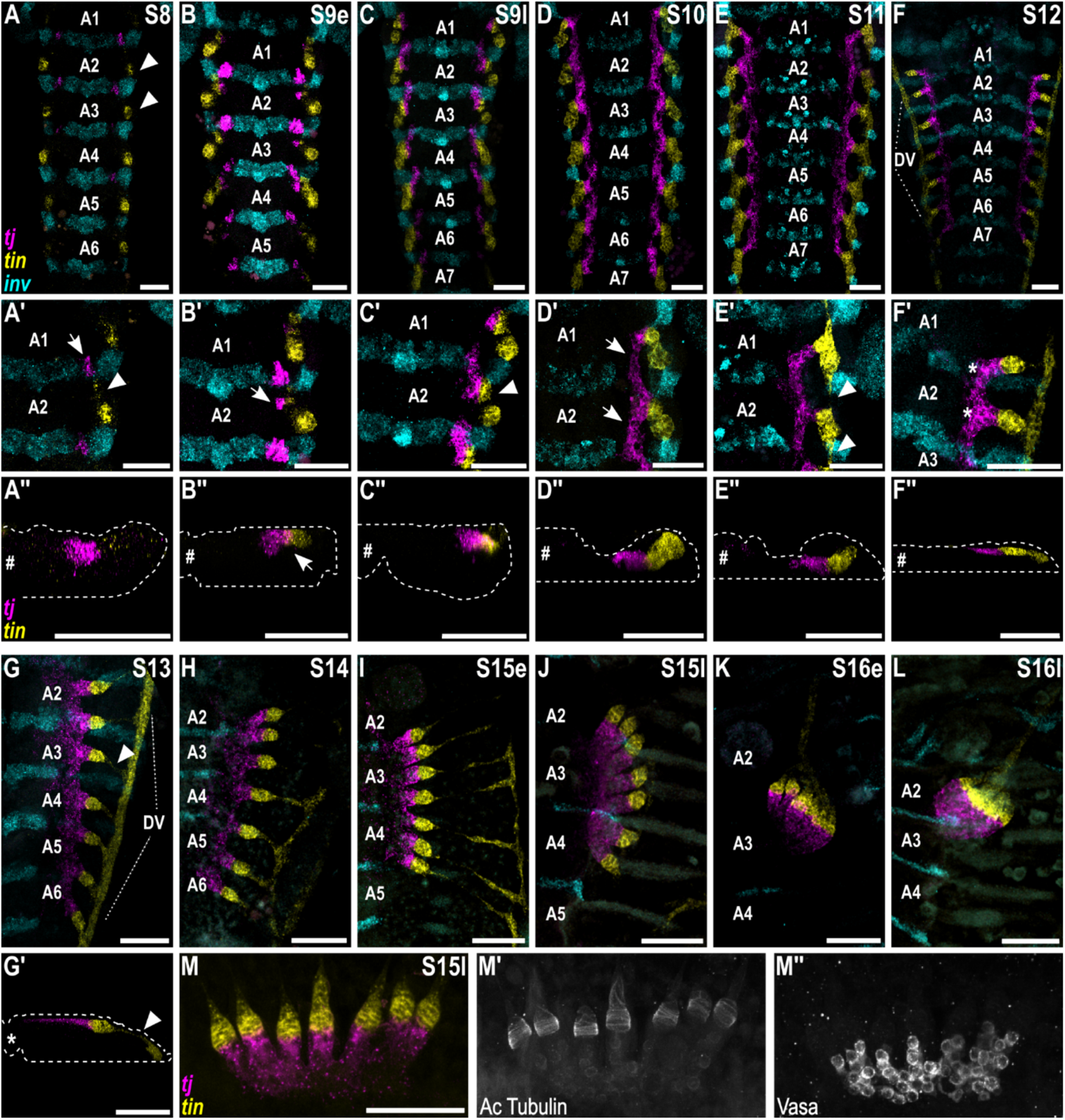
Segment-specific domains of *Ofas*-*traffic jam* and *Ofas*-*tinman* expression form the somatic gonad precursors. **(A-L)** Selected abdominal regions where *Ofas*-*tin* and *Ofas*-*tj* expression are detected across sequential stages of *O. fasciatus* embryonic development. *Ofas-tj* (magenta) and *Ofas-tin* (yellow) mark somatic gonad precursors. *Ofas*-*inv* (cyan) marks the posterior compartment of each segment. Segment identity is indicated in each panel. In (A), arrowheads indicate the primary and lateral *Ofas-tin* domain. In (F-G), the dorsal vessel (DV) is indicated with dotted lines. Tapered regions of *Ofas-tin* expression are highlighted with arrowheads in (G). **(A’-F’)** Magnified views of the right hemisegment of segments A1 and A2 from the embryos shown in (A-F). In (A’), arrowhead shows the secondary domain of *Ofas-tin* expression while the arrow points out the primary *Ofas*-*tj* expression domain. In (B’), the secondary domain of *Ofas*-*tj* is highlighted with a white arrow. In (C’), the adjacent *Ofas-tin*/*Ofas*-*tj* units are indicated with a white arrowhead. White arrows and arrowheads in (D’) and (E’) show the expansion of the *Ofas*-*tj* and *Ofas-tin* domains leading the formation of a contiguous expression domain from A1-A7. Asterisks in (F’) show individual *Ofas-tin*/*Ofas*-*tj* gonadiole precursors. **(A’’-F’’ and G’)** Transverse sections through the right hemisegment of segment A2. Only *Ofas-tin* and *Ofas*-*tj* are visualized in these panels. Dotted lines indicate the embryonic surface while the hash denotes the midline. In (B’’), the arrowhead shows the adjacent expression of *Ofas-tin* and *Ofas*-*tj*. In (G’), the arrowhead highlights the tapering *Ofas-tin* domain. **(M-M’’)** show a coalesced embryonic gonad in a late-stage 15 embryo. *Ofas*-Vas labels the germ cells which are present in the *Ofas*-*tj* -positive (magenta) region of the gonad. Acetylated Tubulin (Ac Tubulin) is detected in the *Ofas-tin* - positive (yellow) region of the gonad and at lower levels in the germ cells. Scale bars = 50 µm.

We first detected *Ofas*-*tin* expression at stage 8 in segmentally repeated domains along the lateral margins of the mesoderm (Figure 2A; white arrowheads), coincident with the formation of coelomic cavities. These primary expression domains appeared sequentially from anterior to posterior, beginning in the labial segment (Figure S2A, A′; white arrowheads). During stage 8, a second *Ofas*-*tin*-expressing domain emerged sequentially only in abdominal segments A1–A7 (Figure 2A′; white arrowhead), positioned anterior and medial to the primary *Ofas*-*tin* domain within each segment. At the same stage, we detected mesodermal *Ofas*-*tj* expression appearing sequentially in the posterior compartments based on co-expression with the *O. fasciatus* ortholog of *invected* (*Ofas-inv*) ^72^ of segments A1–A6 (Figure 2A′; white arrow), but not in the head or thoracic segments (Figure S2A, A″). By early stage 9, secondary *Ofas*-*tj*-expressing domains emerged immediately adjacent to the secondary *Ofas*-*tin* domains (Figure 2B, B′–B″; white arrows). By the end of stage 9, the initially separated medial secondary *Ofas*-*tj*- and lateral *Ofas*-*tin*-expressing domains formed seven repeated *Ofas*-*tin* and *Ofas*-*tj* units along the anteroposterior (AP) axis in segments A1–A7 (Figure 2C, C′; white arrowhead).

At stage 10, the *Ofas*-*tj*-expressing domains in A2–A7 expanded anteriorly and appeared contiguous along the AP axis (Figure 2D–D′; white arrows). In contrast, the A1 *Ofas*-*tj* domain was not comparably anteriorly expanded (Figure 2D–D′). At stage 11, the secondary *Ofas*-*tin*-expressing domains also appeared expanded and were connected with each other, resulting in contiguous *Ofas*-*tin*- and *Ofas*-*tj*-expressing domains associated with seven repeated structures, one in each of segments A1 to A7 (Figure 2E, E′; white arrowheads).

At stage 12, we detected *Ofas*-*tin* expression along each lateral side of the embryo from the labial segment to the posterior abdomen (Figure 2F; DV). We refer to these as longitudinal *Ofas-tin*-expressing domains, and interpret them as being derived from the primary *Ofas-tin* domains first observed at stage 8 (Figure 2A; white arrowheads). We speculate that these longitudinal *Ofas*-*tin*-expressing domains may correspond to the developing dorsal vessel (Figure 2F–G; DV). In addition, seven tube-like *Ofas*-*tin*-expressing structures, corresponding spatially to the earlier secondary *Ofas*-*tin* domains, extended medially from the longitudinal *tin*-expressing domains (Figure 2F, F′; asterisks).

During stages 13–14, as the embryo further shortened and widened, the lateral ends of the tube-like *Ofas*-*tin* expression domains became tapered while remaining connected laterally to the longitudinal *Ofas*-*tin*-expressing domain (Figure 2G–I; white arrowheads in G–G′). We interpret these tube-like *Ofas*-*tin*-expressing domains as gonadiole primordia, noting that they are associated with PGCs and also correspond spatially to the region enriched in acetylated Tubulin in each segment (Figure 2M–M″). We therefore considered this connected group of tube-like *Ofas*-*tin* and *Ofas*-*tj*-expressing cells to be the embryonic gonad primordium.

The embryonic gonad primordium initially formed in the dorsal mesoderm of the embryo (Figure 2A’’–F’’) and extended across abdominal segments A1–A7 up to stage 12 (Figure 2F). Following katatrepsis (stage 13) and during dorsal closure (stage 14 onward), it progressively condensed along the AP axis (Figure 2G–J). During this process, the lateral ends of the *Ofas*-*tin*-positive structures appeared to shift anteriorly and converge (Figure 2K). The morphology and final position of these structures are consistent with their contribution to the previously described suspensory ligaments that connect the gonads to the thoracic body wall ^61^. Concurrently, the seven gonadiole primordia became bundled and reoriented approximately parallel to the AP axis (Figure 2K). Nearly two days prior to hatching (stage 16l), each embryonic gonad had condensed to lie almost entirely within segment A2, excluding the suspensory ligaments and primordial gonoduct (Figure 2L). We detected no consistent differences in morphology or in *Ofas*-*tin* or *Ofas*-*tj* expression that were suggestive of sexual dimorphism from stage 8 through late stage 16 (nearly two days before hatching), consistent with previous reports that sexual differentiation of the gonads is not apparent during these embryonic stages ^73^.

### Ofas-tin and Ofas-tj *expression patterns are mutually independent in gonadiole-forming domains, and neither is required for gonadiole formation*

Because gonadiole primordia comprise cells associated with both the *Ofas*-*tin* and *Ofas*-*tj* expression domains (Figure 2), we wished to determine whether reduction of one or both genes would disrupt gonadiole primordia formation. We therefore performed embryonic RNA interference (eRNAi) targeting *Ofas*-*tin* and *Ofas*-*tj* individually (Figure 3A–C″) or simultaneously (Figure 3D–G″), and confirmed reduction of target transcript with hybridisation chain reaction (HCR) (Figure S3). These experiments yielded two major results relevant to gonadiole number determination. First, we found that expression of these two genes in gonadiole primordia was mutually independent. Following *Ofas*-*tin* RNAi, the *Ofas*-*tj* expression domains, organized into seven primordia, remained detectable at stage 13 in all embryos examined (*n* = 135/135; Figures 3B, S6A; Tables S4, S5), and formed primordia comparable to those observed in eRNAi control embryos (*n* = 209/209; Figures 3A, S6A; Tables S4, S5). Similarly, *Ofas*-*tin* expression remained detectable following *Ofas*-*tj* eRNAi (*n* = 41/41; Figures 3C, S6A; Tables S4, S5), organized into seven primordia. In both eRNAi conditions, however, knockdown appeared incomplete, based on faint expression of the targeted gene detected with HCR (Figure 3B′–C′’; white arrowheads). Thus, these results indicate that partial reduction of *Ofas*-*tin* or *Ofas*-*tj* expression does not abolish the expression domain of the other gene, although the incomplete knockdown precludes us from excluding a requirement for *Ofas*-*tin* or *Ofas*-*tj* in establishing or maintaining the respective expression domains.

**Figure 3.**
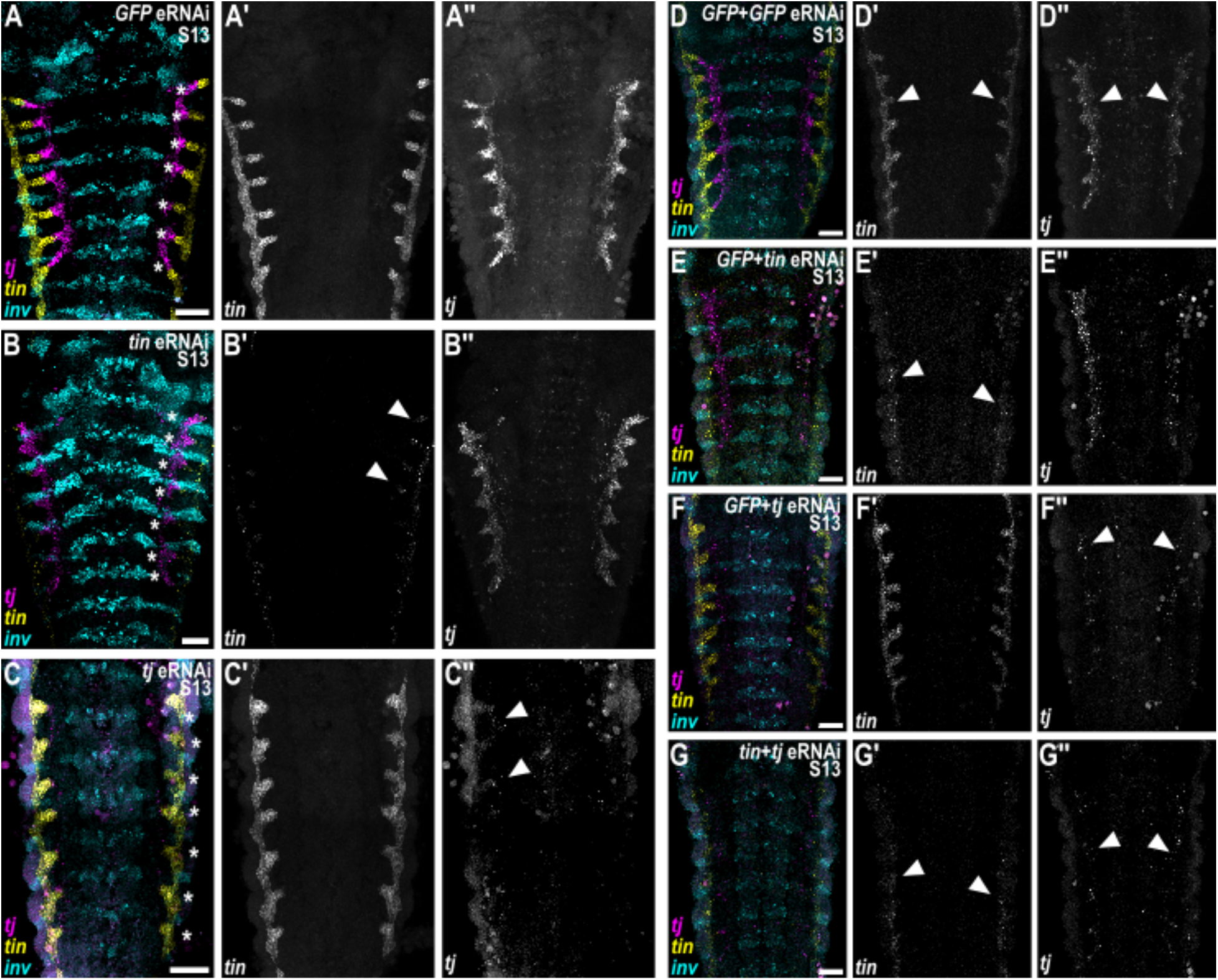
*Ofas*-*tin* and *Ofas*-*tj* are expressed independently of each other. **(A-C’’)** Representative images of selected abdominal segments of embryos from double eRNAi experiments. (A-A’’) A *GFP* dsRNA-injected (*GFP* eRNAi) embryo (n=209/209); (B-B’’) an *Ofas*-*tin* dsRNA-injected (*tin* eRNAi) embryo (n=135/135). *Ofas*-*traffic jam* (*tj*) expression is still detectable, and seven gonadiole primordia still form. *Ofas*-*tin* expression is still present weakly (B’; white arrowheads); (C-C’’) *Ofas*-*tj* dsRNA-injected (*tj* eRNAi) embryo (n=41/41). *Ofas*-*tin* expression does not appear to be affected while *Ofas*-*tj* continues to be weakly expressed (C’’; white arrowheads). **(D-G’’)** Representative images of selected abdominal segments of embryos from double eRNAi experiments. (D-D’’) A *GFP+GFP* double eRNAi (*GFP*+*GFP* eRNAi) embryo (n=14/14); (E-E’’) *GFP*+ *Ofas*-*tin* dsRNA-injected (*GFP*+*tin* eRNAi) embryo (n=17/19). *Ofas*-*traffic jam* (*tj*) expression does not appear to be affected and forms seven precursors (E’’). *Ofas*-tinman (*tin*) expression is still present weakly (E’; white arrowheads); (F-F’’) a *GFP*+ *Ofas*-*tj* -dsRNA-injected (*GFP*+*tj* eRNAi) embryo (n=15/16). *Ofas*-*tin* expression does not appear to be affected (F’) while *Ofas*-*tj* continues to be weakly expressed (F’’; white arrowheads). Seven precursors are formed. (G-G’’) a *tin*+ *tj*-dsRNA-injected (*tin*+*tj* eRNAi) embryo (n=20/23). Weak expression of both tin and tj is detectable (G’-G’’; white arrowheads) and precursors-like structures are visible. *Ofas*-*tj* (magenta) and *Ofas*-*tin* (yellow) mark somatic gonad precursors. *Ofas*-*inv* (cyan) marks the posterior compartment of each segment. Asterisks indicate individual gonadiole primordia. Scale bars = 50 µm.

Second, we observed that in both single knockdown conditions, seven gonadiole primordia remained identifiable (Figure 3E–E″, F–F″; S5). In the double knockdown condition, expression of both genes remained detectable with HCR, albeit at lower levels than in controls, and we could also still detect formed gonadiole structures (Figure 3G–G’’). However, because of the residual expression noted above (Figure 3E’, F”, G’–G″; white arrowheads), we cannot rule out the possibility that complete loss of *Ofas*-*tin* and *Ofas*-*tj* would disrupt gonadiole primordia specification or formation.

### Germ cells populate the gonad after gonadiole primordia formation

The *O. fasciatus* embryonic gonad at stage 15 comprises two cell populations: the somatic gonad precursors and the PGCs (Figure 2M–M”). PGCs are first observed at the cellular blastoderm stage (Table S2; stage 3). We ^21^ and others ^74^ previously described the location of PGC emergence as being immediately inside the posterior pit, which is an indentation at the ventral posterior of the blastoderm into which the germ band invaginates during its formation. PGCs then migrate towards the anterior, along the dorsal side of the germ band ^21,63,74,75^. By stage 8, most PGCs are organised into segmental clusters, with one cluster located in each of abdominal segments A4–A6; many embryos also have an additional cluster in A7 (Figure 4A). At stage 9, coincident with neuroectodermal invagination, each cluster separates bilaterally along the ventral midline, to yield a total of six to eight clusters, one in each hemisegment of A4 through A6 or A7 (Figure 4B–C). During stages 10–12, the clusters on each of the left and right sides of the embryo merge to form two longitudinal groups of PGCs oriented parallel to the AP axis (Figure 4D–F), spanning abdominal segments A4–A6. By stage 13, small numbers of cells from these longitudinal groups begin to appear in columnar formations perpendicular to the AP axis of the embryo (Figure 4G; asterisks). By early stage 15, seven distinct such columnar groups of PGCs are distinguishable (Figure 4H; asterisks). In contrast, the seven somatic gonadiole primordia are already established by late stage 9, including in segments A1–A3 (Figure 2C), where PGCs are not present (Figure 4C). Thus, the seven somatic gonadiole primordia are established before PGCs become organised into seven corresponding groups. This temporal sequence suggests that PGCs are unlikely to determine gonadiole number.

**Figure 4.**
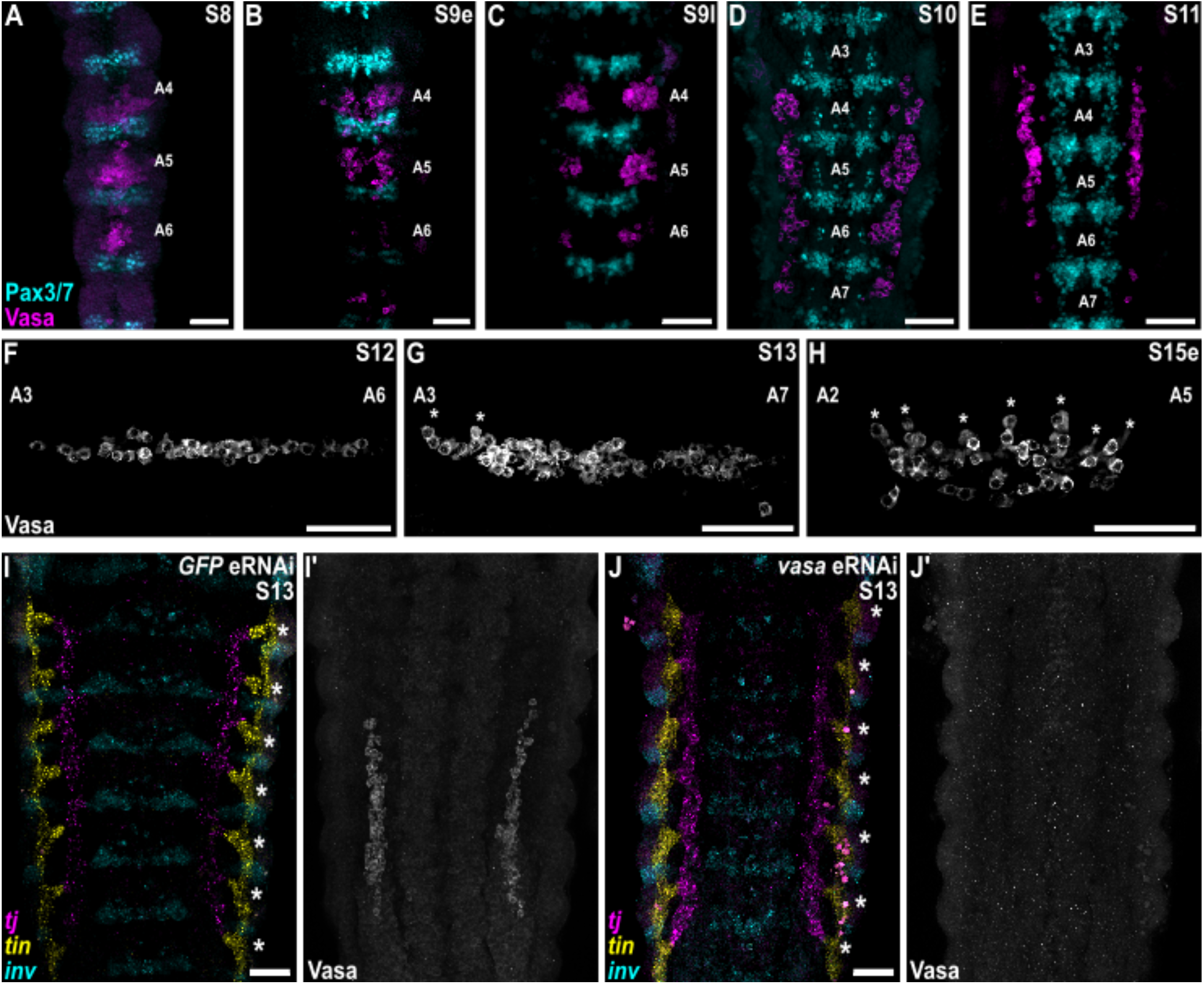
Germ cells begin to form seven independent clusters after the somatic gonad precursors are established. **(A-H)** Germ cells, labelled by Ofas-Vas (magenta in A-E; grey in F-H), across sequential stages of embryonic development. Pax3/7 (cyan) marks the posterior compartment of each segment up to S8 and segmentally iterated neuroblast clusters thereafter. **(I-J’)** Representative images of selected abdominal segments of (I-I’) a *GFP* dsRNA-injected embryo (*GFP* eRNAi, n=49/49); (J-J’) an *Ofas*-*vas* dsRNA-injected embryo (*Ofas*-*vas* eRNAi, n=32/32).

To ask whether PGCs influenced gonadiole morphogenesis or final gonadiole number determination, we used RNAi to abrogate the function of the *O. fasciatus* homolog of the *vasa* gene (*Ofas-vas*), which is required for survival of many ^76,77^ (but not all; see ^22,78^) studied animal germ cells ^24^. *Ofas*-*vas* is expressed throughout embryonic development (Figure S4A), with its transcript detectable at low levels in most embryonic cells, and at somewhat higher levels in PGCs ^21,75^. Ofas-Vas protein is specifically detectable in PGCs during their formation, migration, and gonad coalescence ^63^. We found that depletion of *Ofas*-*vas*, either embryonically (eRNAi; efficacy confirmed by assessment of Ofas-Vas protein expression; Figure S6B; Table S6) or maternally (through depletion in adult parental females, pRNAi; efficacy confirmed by Ofas-Vas protein expression; Figure S6B; Table S6), did not appear to alter PGC organisation (Figures 4J–J′, S4C–C′), and did not change gonadiole number (eRNAi *n* = 32/32; pRNAi *n* = 12/12; Figures 4J–J’, S4C–C’, S6A; Tables S4, S7). Though we did not detect Ofas-Vas expression in *Ofas-vas* RNAi embryos (Figures 4J’, S4C,S6B; Table S6), we were nonetheless able to identify PGCs based on their low levels of acetylated Tubulin expression (Figure S4C-C’). *Ofas*-*vas* function therefore appears dispensable for the correct establishment of seven gonadiole primordia. However, because, as previously reported ^79^, PGCs were not eliminated by *Ofas*-*vas* RNAi (Figure S4C–C’), we cannot formally exclude a role for PGCs in this process.

### Reduction of gonadiole-producing segments leads to fewer gonadiole primordia

Our observation of *Ofas*-*tin* and *Ofas*-*tj* expression in segmentally repeated groups of somatic gonadiole primordia suggested that the number of gonadiole-producing segments might determine the number of gonadioles formed. We therefore hypothesised that reducing the number of these segments would reduce gonadiole primordium number. To test this hypothesis, we examined somatic gonad development following two distinct perturbations of the segmentation gene regulatory network.

First, we knocked down the *O. fasciatus* homolog of the gap gene *giant* (*gt*) ^80^. Gap genes are expressed in and required for the specification of contiguous regions of the insect body plan, and disruption of their function can result in the loss of groups of adjacent segments ^81–83^. In *O. fasciatus*, the first four abdominal segments (A1–A4) are missing in *Ofas-gt* knockdown embryos ^80^. Because A1–A4 normally contribute four of the seven segmentally repeated gonadiole primordia, we predicted that their loss would reduce the number of gonadiole primordia. Consistent with this prediction, stage 13 *Ofas*-*gt* dsRNA-injected embryos (eRNAi) showed both reduced numbers of abdominal segments (Figures 5B–C, S6B; Table S6) as previously reported ^80^, and a reduction in gonadiole primordium number that corresponded to the reduced number of abdominal segments (*n* = 40/91; Figures 5A–C, S6A; Tables S4, S5).

**Figure 5.**
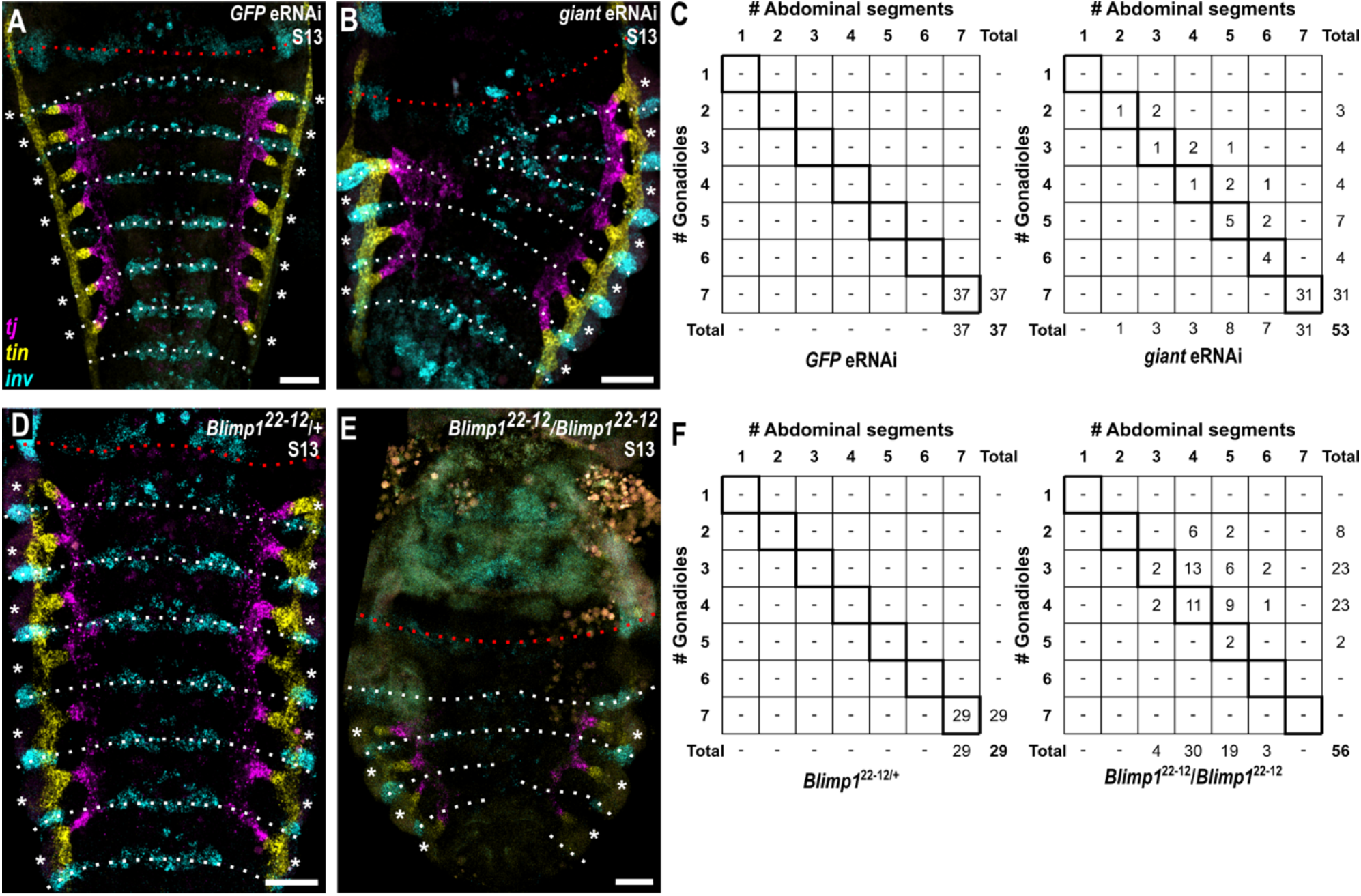
Reducing the number of gonadiole-producing segments decreases gonadiole number. **(A-B, D-E)** Representative images of selected abdominal segments of embryos (A) injected with dsRNA targeting *GFP* (*GFP* eRNAi, n=66/66); (B) injected with dsRNA targeting *Ofas-giant* (n=40/91 show a reduction in gonadiole primordium number); (D) *Ofas*-*Blimp1*^22–12^*/+* heterozygotes (n=29/29) have wild type numbers of gonadiole precursors; (E) *Ofas*-*Blimp1*^22–12^ homozygotes (n=56/56) show a reduction in gonadiole primordium number. *Ofas-tj* (magenta) and *Ofas*-*tin* (yellow) mark somatic gonad precursors. *Ofas*-*inv* (cyan) marks the posterior compartment of each segment. Red dotted line denotes the posterior of the third thoracic segment and white dotted lines delineate the seven abdominal segments. Asterisks indicate individual gonadiole primordia. Scale bars = 50 µm. **(C, F)** Plots showing the relationship between gonadiole precursor number (y-axis) and the number of the first seven abdominal segments still present (x-axis) in (C) *GFP* eRNAi and *Ofas*-*giant* eRNAi embryos and (F) *Ofas*-*Blimp122–12*heterozygous and *Ofas*-*Blimp1^22–12^homozygous* mutant embryos in each hemisegment. The diagonal indicates a perfect one-to-one correspondence between the total number of gonadiole primordium-producing segments and gonadiole primordia number in each hemisegment.

Second, we examined gonadiole primordium number in *O. fasciatus* mutants homozygous for a previously generated hypomorphic *Ofas-Blimp1*^22–12^ allele, which is predicted to yield a non-functional protein product missing the zinc finger domains ^84^. In embryos of these mutants, the segment pairs T1/T2, T3/A1, A2/A3, A4/A5 and A7–A10 are consistently fused, while A6 is sometimes unaffected, and sometimes incorporated into the A7–A10 fusion ^84^. We found that at stage 13, homozygous *Ofas*-*Blimp1* mutant embryos contained only two to four gonadiole primordia (*n* = 56/56; Figures 5D–F, S6A; Table S4), and had reduced numbers of abdominal segments (Figures 5E-F, S6B; Table S6) as previously reported ^84^. The relationship between the number of morphologically distinguishable segments and gonadiole number was less pronounced in *Ofas*-*Blimp1* mutants (Figure 5F) than following *Ofas-gt* knockdown (Figure 5C). One possible explanation is that fusion between adjacent segments alters or produces variable segmental identities, such that the number of morphologically distinguishable segments does not directly reflect the number or identity of segmental domains competent to produce gonadiole primordia. Together these findings support a model in which the segmental organisation of the embryo establishes the repeated somatic gonadal units that determine the final number of gonadioles. Thus, loss of any of segments A1–A7 results in correspondingly fewer gonadiole primordia.

### Posterior Hox genes regulate gonadiole primordium formation

Our observations of wild type embryos and of embryos with abnormal numbers of abdominal segments suggested that the segmental organisation of the embryo establishes the repeated somatic gonadal units that determine the final number of gonadioles. Further, the restriction of gonadiole primordia to A1–A7 suggested that segmental identity may determine whether a segment is competent to form a gonadiole primordium. The conserved role of Hox genes in specifying regional identity along the AP axis of animals ^85^ led us to hypothesise that they might be involved in conferring this competence upon some, but not all, abdominal segments in *O. fasciatus*. Previous work showed that, the posterior Hox genes *Ultrabithorax* (*Ofas*-*Ubx*), *abdominal A* (*Ofas*-*abd-A*) and *Abdominal B* (*Ofas*-*Abd-B*) pattern the *O. fasciatus* abdomen ^86^, and we detected their expression in the abdominal mesoderm before and during gonadiole primordium formation (Figures 6A, S5A–B). We therefore asked whether these genes regulate the segment-specific formation of gonadiole primordia.

**Figure 6.**
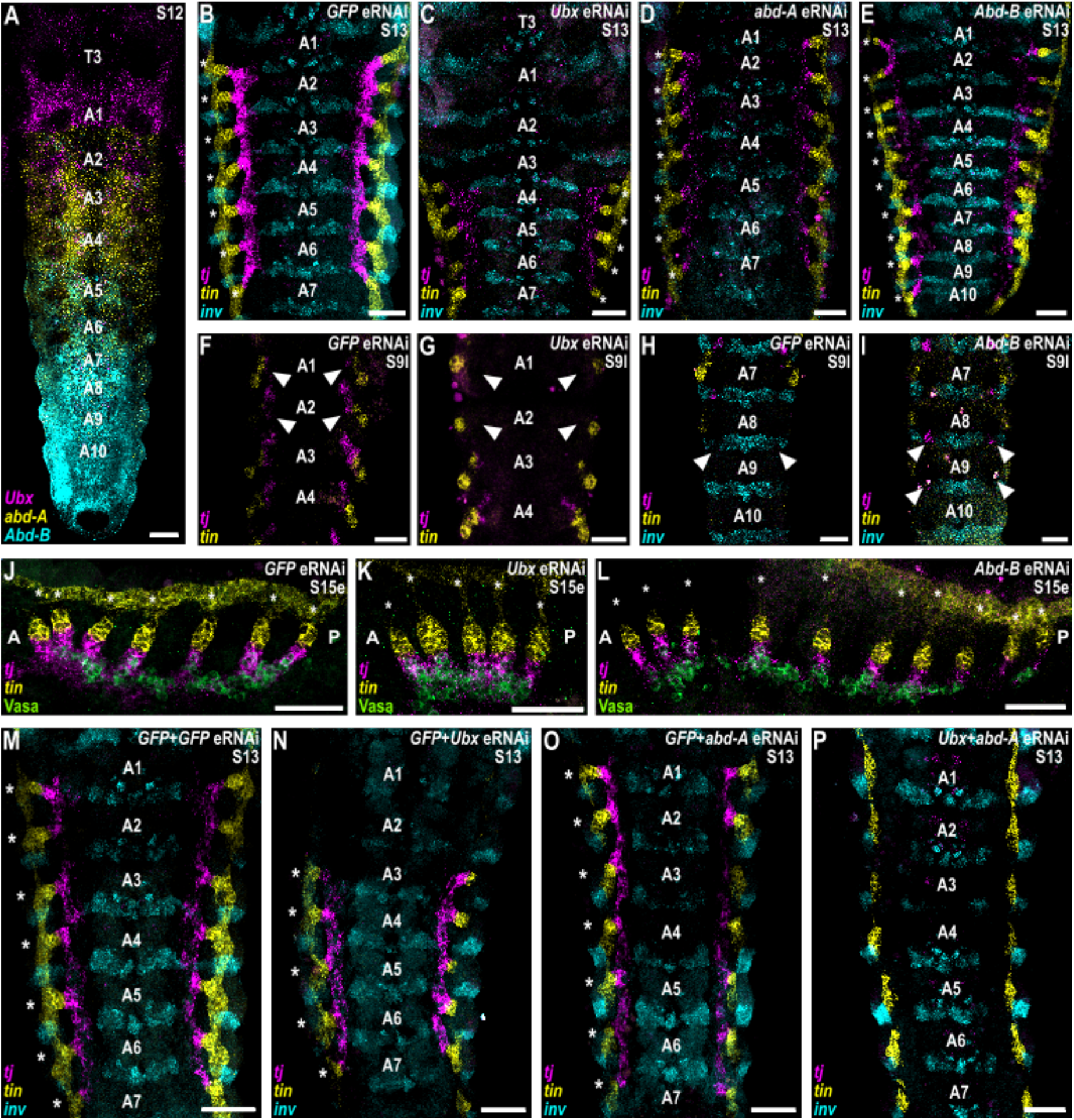
Posterior Hox genes regulate gonadiole primordia formation. **(A)** Posterior thoracic and abdominal expression of *Ofas*-*Ultrabithorax* (*Ubx*, magenta), *Ofas*-*abdominal A* (*abd-A*, yellow) and *Ofas*-*Abdominal B* (*abd-B*, cyan) at stage 12. Individual expression patterns are shown in Figure S5A. Scale bar = 50 µm. **(B-E)** Representative images of selected abdominal segments of embryos injected with dsRNA targeting (B) *GFP* (*GFP* eRNAi), (C) *Ofas*-*Ubx*, (D) *Ofas*- *abd-A* or (E) *Ofas*-*Abd-B*. *Ofas*-*Ubx* eRNAi embryos form only five gonadiole precursors, with the precursors missing in A1 and A2 (n=34/56). *Ofas*-*Abd-B* eRNAi embryos form ectopic gonadiole precursors in A8-A10 (n=32/60). *GFP* eRNAi (n=69/69) and *Ofas*-*abd-A* eRNAi (86/87) embryos show no change in gonadiole precursor number. **(F-G)** Neither *Ofas*-*tj*-nor *Ofas*-*tin*-expressing somatic gonad precursors are specified at stage 9 in A1 or A2 in *Ofas*-*Ubx* eRNAi embryos (white arrows). **(H-I)** In *Abd-B* eRNAi embryos, ectopic *Ofas*-*tj*- and *Ofas*-*tin*-expressing somatic gonad precursors are specified in segments A8-A10 in stage 9 in the same location within each segment as in the anterior abdominal segments (white arrows). **(J-L)** In *Ofas*-*Ubx* and *Ofas*-*Abd-B* eRNAi embryos, gonads coalesce normally despite having an abnormal number of gonadioles. Anterior (A) and posterior (P) are indicated on the images. **(M-P)** Representative images of selected abdominal segments of embryos from double eRNAi experiments. *Ofas*-*Ubx*+ *Ofas*-*abd-A* double eRNAi embryos have no detectable gonadiole precursors in any segment (n=102/112). Lateral secondary *Ofas*-*tin* domains connect to each other forming a contiguous expression domain. *GFP*+ *GFP* double eRNAi (*GFP*+*GFP* eRNAi n=98/98) and *GFP*+ *Ofas*-*abd-A* double eRNAi (n=79/79) embryos show no change in gonadiole precursor number, whereas *GFP*+ *Ofas*-*Ubx* double eRNAi embryos have only five precursors (n=89/102). In panels (B-P), *Ofas*-*tj* is shown in magenta, *Ofas*-*tin* in yellow, *Ofas*-*inv* in cyan, and *Ofas*-Vas in green. Asterisks denote individual gonadiole primordia. Scale bars = 50 µm.

We individually depleted *Ofas*-*Ubx*, *Ofas*-*abd-A* and *Ofas*-*Abd-B* using eRNAi. We confirmed the efficacy of eRNAi based on homeotic transformations observed in the RNAi embryos (Figure S6B; Table S6). In *Ofas*-*Ubx* knockdown embryos, the secondary *Ofas*-*tin* and both *Ofas*-*tj* domains were not detected at early stage 9 in A1 or A2 (Figure 6G), and gonadiole primordia were also absent from A1 and A2 at stage 13, resulting in five gonadiole primordia (*n* = 34/56; Figures 6C, S6A; Tables S4, S5). In contrast, *Ofas*-*Abd-B* knockdown resulted in ectopic *Ofas-tin* and *Ofas-tj* expression domains in A8 and A9 at stage 9 (Figure 6I), and ectopic gonadiole primordia in A8–A10 at stage 13, producing a total of ten gonadiole primordia (*n* = 23/60; Figures 6E, S6A; Tables S4). Despite their abnormal gonadiole numbers, however, the primordial gonads of *Ofas*-*Ubx* and *Ofas*-*Abd-B* eRNAi embryos underwent normal coalescence (Figure 6J–L).

Depletion of *Ofas*-*abd-A* alone had little apparent effect on gonadiole primordium formation, with seven gonadiole primordia observed in nearly all embryos (*n* = 86/87; Figures 6D, S6A; Tables S4). A previous study provided evidence that *Ofas-abd-A* knockdown leads to expansion of *Ofas-Ubx* expression domain into posterior abdominal segments, a phenomenon known as posterior prevalence ^87–90^ This results in formation of ectopic legs (which are normally only formed in the thorax) in abdominal segments A2–A8 ^86^. We therefore hypothesised that *Ofas-abd-A* RNAi embryos generate gonadiole primordia in A2–A7 because *Ofas-Ubx* expression expands posteriorly and promotes their development. This hypothesis predicts that *Ofas*-*abd-A* and *Ofas*-*Ubx* might play overlapping or redundant functions in promoting gonadiole primordium formation. To test this prediction, we performed double eRNAi for *Ofas*-*Ubx* and *Ofas*-*abd-A* simultaneously. In contrast to either single knockdown, combined depletion of these two genes resulted in complete loss of the secondary *Ofas*-*tin* and *Ofas*-*tj* expression domains at stage 11 (Figure S5O–R), and loss of gonadiole primordia at stage 13 (*n* = 102/112; Figures 6M–P, S6A; Tables S4). However, the longitudinal *Ofas*-*tin* expression domain that likely corresponds to the developing dorsal vessel appeared unaffected (Figures 6M–P, S5O–R; Tables S4). We conclude that *Ofas*-*Ubx* and *Ofas*-*abd-A* are specifically required for the *Ofas*-*tin* and *Ofas*-*tj*-expressing domains associated with gonadiole primordium formation, but not for the primary *Ofas*-*tin*-expressing domain.

The results of parental RNAi (pRNAi) experiments were consistent with those of the embryonic RNAi experiments with respect to the segments in which gonadiole primordium formation was affected (Figures S5C–F, S6A; Table S4, S7). In nymphs produced by injected mothers, we observed morphological defects consistent with the homeotic transformations reported in previous studies of Hox genes using pRNAi (Figure S5G–L) ^86^. These transformations were weaker than those observed with eRNAi (Figures S5C–F, S6A-B; Tables S4 S6). Accordingly, for example, in A1 of *Ofas-Ubx* pRNAi embryos, we observed gonadiole primordia that appeared incomplete, or disconnected from the rest of the primordia at stages when control embryos had all seven primordia linked through their *Ofas-tj* domain (*n* = 25/25 had seven gonadioles that were at least partially formed, but 11/25 of these were disconnected in A1; Figures S5D, S6A; Table S4). Similarly, in A8 and A9 of *Ofas-Abd-B* pRNAi embryos, we observed gonadiole primordia in 5/16 embryos, but in all of these embryos, the gonadiole primordia appeared incomplete (Figures S5F, S6A; Table S4).

The *O. fasciatus* ortholog of the Hox gene *Antennapedia* (*Ofas*-*Antp*) is expressed predominantly in the thoracic segments immediately anterior to the *Ofas-Ubx domain* ^86^. Based on the principle of posterior prevalence, *Ofas-Antp* expression is predicted to expand into segments A1–A2 upon *Ofas-Ubx* knockdown. Thus, the loss of gonadiole primordia from A1–A2 in the *Ofas-Ubx* condition (Figure 6C, G) could result either from the loss of a primordium-promoting function of *Ofas-Ubx* in A1–A2, or to the gain of a primordium-suppressing function of *Ofas-Antp* in those segments. To try to distinguish between these alternatives, we knocked down *Ofas-Antp* with eRNAi. We found that depletion of *Ofas-Antp* did not affect gonadiole primordium formation, either alone (*n* = 35/35; Figures S5M-N, S6A; Tables S4), or in combination with *Ofas-Ubx* knockdown (*n* = 7/42 had seven gonadioles, while n = 35/42 had five gonadioles, recapitulating the *Ofas-Ubx* single eRNAi phenotype; Figures S5S–V, S6A; Table S4). These data suggest that loss of *Ofas-Antp* activity is not sufficient to confer gonadiole-forming identity upon thoracic segments.

## Discussion

We have shown that unlike in *Drosophila*, where adult ovariole number is determined through larval proliferation and terminal filament assembly, the number of gonadioles in *O. fasciatus* is established through the segmental specification of discrete primordia during embryogenesis. Our results further suggest that posterior Hox genes determine the abdominal segments that are competent to produce gonadioles. Under this model, one or both of *Ofas*-*Ubx* and *Ofas*-*abd-A* are required for formation of the secondary *Ofas*-*tin* and *Ofas*-*tj* expression domains in A1–A7 that prefigure gonadiole primordium formation, and *Ofas*-*Abd-B* restricts gonadiole formation posteriorly, either by repressing *Ofas-tin* and *Ofas-tj* transcription in these domains, or by failing to promote it.

### Hox genes in somatic gonad precursor specification

Our Hox gene knockdown experiments suggest that *Ofas*-*Ubx* and *Ofas*-*abd-A* promote gonadiole primordium formation in their respective expression domains (Figure 6), suggesting that they regulate the same target genes in these positions. There is ample precedent for such a regulatory paradigm: these two Hox genes can occupy common binding sites ^91^, and share regulatory targets during body-wall and alary muscle determination ^92,93^ and appendage formation ^91^ in *D. melanogaster*, and abdominal trachea formation of *O. fasciatus* ^94^.

As we report here for *O. fasciatus*, somatic gonad precursor (SGP) specification in *D. melanogaster* also depends on the Hox genes *abd-A* and *Abd-B*. In *D. melanogaster*, Abd-A alone specifies an anterior SGP population in parasegments 10 and 11, whereas Abd-A and Abd-B jointly specify a posterior SGP population in parasegment 12 ^9^. These SGP populations are molecularly distinct: *escargot* (*esg*) is expressed only in the anterior population, whereas *eyes-absent* (*eya*) is expressed only in the posterior population ^9^. In the adult ovary, *esg* is expressed in escort cells ^95^, whereas *eya* is expressed in follicle stem cells ^96^, suggesting the possibility that the two embryonic cell populations might give rise to distinct cell types in *D. melanogaster* adult ovaries. In *O. fasciatus*, however, we observed no morphological differences between adult gonadioles whose specification required *Ofas*-*Ubx* (A1–A2) and those specified by *Ofas*-*abd-A* alone (A3–A7).

Formation of both primary and secondary *Ofas*-*tj*-expressing domains depends on *Ofas*-*Ubx* or *Ofas*-*abd-A*, whereas the two *Ofas*-*tin*-expressing domains are differentially regulated - Following *Ofas*-*Ubx* and *Ofas*-*abd-A* double knockdown, which transforms the affected segments towards more anterior identities, the primary *Ofas*-*tin* domains still form and give rise to the presumptive dorsal vessel (Figures 6P, S5R). However, the secondary *Ofas*-*tin* domains, which contribute to the somatic gonadiole precursors, fail to form (Figures 6P, S5R). These results suggest that *Ofas*-*Ubx* and *Ofas*-*abd-A* are specifically required for the formation of the somatic gonad-associated *Ofas*-*tj* and secondary *Ofas*-*tin* domains, but not for the primary *Ofas*-*tin* domain that is putatively associated with the dorsal vessel.

We find that depletion of either *Ofas-tin* or *Ofas-tj* does not abolish expression of the other gene, and that gonadiole precursors still form even if both are depleted. This could be because we were unable to achieve complete knockdown, such that the low levels of residual *Ofas-tin* and *Ofas-tj* expression remaining were sufficient to support gonadiole precursor formation. Alternatively, these genes may not be required for gonadiole precursor specification, but may instead act following specification to regulate the morphogenesis or function of these cells. This possibility is consistent with findings in *D. melanogaster*, where *traffic jam* is not required for SGP specification ^70^, but is expressed in SGPs as they come into contact with PGCs ^70,97^, and is required for intermingling of somatic and germ cells during embryogenesis ^70^. Similarly, *raw*, a negatively regulator of JNK signaling, is expressed in SGPs and required for the extension of SGP protrusions that surround the germ cells during gonad coalescence, but is dispensable for SGP specification ^98^.

The initial mesodermal domains of *Ofas*-*tj* expression in segments A1–A7 directly overlie ectodermal stripes of *Ofas*-*inv* expression (Figure 2A–B), which also express *Ofas*-*hedgehog* ^99^. Similarly, somatic gonad precursors in *D. melanogaster* arise from mesoderm overlying ectodermal stripes of *engrailed* and *hedgehog* expression ^100^. Future studies should investigate the hypothesis that Hedgehog signaling contributes to the specification of *tj*-expressing somatic gonad precursors in *O. fasciatus*, as it does in *D. melanogaster* ^100^.

By contrast, no mesodermal expression domain analogous to the secondary *Ofas*-*tin* domain that produces *Ofas*-*tin*-expressing somatic gonad precursors has, to our knowledge, been described in *D. melanogaster*. Future studies could also test the role of Hedgehog signalling in specifying the *Ofas*-*tj*-expressing population and identify the signals that specify the *Ofas*-*tin*-expressing population. Such experiments, however, will likely require spatial or temporal control of gene knockdown or overexpression to minimize the pleiotropic effects of manipulating Hedgehog and other candidate pathways ^101,102^. To our knowledge, such methods have not yet been established for *O. fasciatus* (but see ^103^). These studies should clarify how the genetic pathways specifying distinct mesodermal cell types have evolved across insects.

### Adult gonad development and function

Wick and Bonhag ^60^ identified four distinct regions of each ovariole in first-instar *O. fasciatus* nymphs: the anterior ovarian strand, posterior ovarian strand, epithelial sheath, and germ cells. Comparison of our embryonic expression data (Figure 2M–M″) with their descriptions leads us to hypothesise that the anterior ovarian strand corresponds to the *Ofas-tin*-expressing domain, whereas the posterior ovarian strand corresponds to the *Ofas*-*tj*-expressing domain.

The anterior ovarian strand ultimately forms the terminal filament, which in *O. fasciatus* is reported to be a syncytial structure at the anterior tip of each adult ovariole ^60,61^. The embryonic *Ofas*-*tin*-expressing domain that gives rise to the terminal filament is directly associated with the dorsal vessel (Figure 2G′) and is enriched in acetylated Tubulin (Figures 1E, 2M’), a marker of stable microtubules ^64,65^. Together with the observation that the *Ofas-tin* expressing domain elongates and becomes more filamentous as development progresses (Figure 2E’’, G’), these features suggest the possibility that the anterior ovarian strand may be subject to mechanical tension. We speculate that during dorsal closure, the cardioblasts migrate and coalesce medially ^104^, thereby exerting force on the terminal filaments. Testing this model in future studies will require direct analysis of tissue movements and mechanical interactions between the dorsal vessel and developing ovary.

The testes and ovaries of first-instar *O. fasciatus* differ in morphology. In the testis, germ cells surround an apical complex that is thought to provide a niche for germline stem cells in adults ^105^. We found that the apical complex expresses *Ofas*-*tin* in adult testioles (Figure S2G–H’), consistent with the possibility that it is derived from the same embryonic cell population as the anterior ovarian strand. Alternatively, adult *Ofas*-*tin* expression may be unrelated to its embryonic expression, and the apical complex may arise from another, unidentified cell population. We also detected *Ofas*-*tj* and *Ofas*-*tin* in regions of the adult testis that house primary spermatogonia and secondary spermatocytes, respectively (Figure S2H–J′). The specific functions of these genes during spermatogenesis, if any, remain unknown.

*Ofas-tj* is also expressed in follicle cells in the adult ovary (Figure S2B–F’). In *D. melanogaster*, *tj* is expressed in intermingled cells, and loss of *tj* disrupts germ-cell ensheathment and differentiation ^70^. The adult expression pattern of *Ofas*-*tj* in *O. fasciatus* suggests that it may perform a similar function in gonadal somatic cells. Furthermore, the failure in oogenesis observed following *Ofas*-*tj* knockdown suggests an additional role in this process (Figure S3A–C).

### Developmental basis of gonadiole number across insects

Since gonadioles are the functional subunits of the adult gonad, their number can constrain an individual’s maximum capacity for gamete production ^8,106^. Here we have provided evidence that the invariant gonadiole number of *O. fasciatus* is determined by the spatial distribution of the specified somatic gonadal precursors, rather than by the PGCs. Similarly, in *Drosophila*, variable ovariole numbers are determined by the number, specification and organization of somatic cells, in this case called terminal filament cells ^18,19,38,41,55^. Thus, in both systems, irrespective of the variability or lack thereof in gonadiole number, this potentially significant component of reproductive fitness appears to be regulated by cellular behaviours within the developing somatic gonad rather than the germ line itself.

Most Heteroptera have an invariant number of seven gonadioles per gonad ^8,107^, and ancestral reconstruction studies have suggested seven as the ancestral gonadiole number for true bugs ^107^. We hypothesise that the developmental and genetic mechanisms establishing this number in *O. fasciatus* may be conserved across the suborder. However, unlike *O. fasciatus*, embryos of the European firebug *P. apterus* are reported to form eight gonadiole primordia, with the anteriormost primordium subsequently degenerating before hatching ^59^. In embryos of the kissing bug *Rhodnius prolixus*, Heming and Huebner ^108^ reported that seven gonadiole primordia form as in *O. fasciatus*, but across segments A3–A6. We hypothesise that evolutionary changes in the Hox-mediated regulation of segmental identity within the Heteroptera could determine segment competence to form, maintain or eliminate SGPs and gonadiole primordia, while still maintaining invariance in adult gonadiole number. This hypothesis is consistent with another instance of Hox-mediated morphological invariance: the case of the number of cervical vertebrae in mammals. In this case, almost all studied mammals have exactly seven vertebrae in the neck ^109–111^, a phenomenon attributed by some authors to Hox-mediated identity regulation of these vertebrae ^110,but^ ^see^ ^112,113^. In sloths, which are one of the two reported exceptions to this rule ^114,115^ (the other being manatees ^109,111,116–118^), a correlative study suggests that changes in Hox gene expression boundaries may promote formation, maintenance or elimination of cervical vertebrae, while maintaining their number invariant within a lineage ^119^.

Invariant ovariole number evolved independently in multiple insect orders ^8^. In the firebrat *Thermobia domestica* (Zygentoma), Woodland ^57^ reported that an invariant number of segmentally organized gonad rudiments first appear during the third nymphal instar. Whether similar mechanisms operate in other lineages with invariant ovariole numbers, such as butterflies and moths (Lepidoptera), which typically have four ovarioles per ovary, or weevils (Curculionoidea: Coleoptera), which typically have two, remains unknown ^8,120^. Notably, to our knowledge no insect species with an invariant ovariole number has been reported to possess more than seven ovarioles per ovary ^8^. Thus, we speculate that the maximum observed invariant ovariole number could potentially be constrained by segmental mechanisms within the 10–12 abdominal segments that characterise the insect body plan.

### Evolutionary implications of invariant morphologies

The timing of gonadiole number specification may influence its sensitivity to environmental conditions, including whether it displays phenotypic plasticity. In *D. melanogaster*, ovariole number is variable, established during larval development ^55,56^, and can display phenotypic plasticity in response to non-genetic factors including nutrition and temperature ^38,41,48,121,122^. The larval stage is a feeding stage, and variation in nutritional input is sensed by larval tissues via the insulin and TOR signaling pathways ^123–127^. The activity levels of these pathways regulate proliferation of most cell types ^128–130^, including somatic gonad cells ^31^. This in turn alters the number of terminal filament cells, the number of terminal filaments formed, and ultimately the number of ovarioles in the adult ^31^. By contrast, gonadiole number in *O. fasciatus* is invariant ^60,61^, and specified during embryogenesis, a non-feeding embryonic stage that by definition is not sensing nutritional inputs. Even if *O. fasciatus* mothers were to experience different nutritional regimes, we predict that the gonadiole number of their offspring would be unaffected, since that number is tied to the Hox code that establishes the invariant body plan. The offspring of starved females can be smaller than those of well-fed mothers, but they still have the same body plan for their species ^121,131,132^. Comparative developmental genetic studies across a broader range of species with invariant and variable gonadiole numbers will be needed going forward, to test the correlation between gonadiole number variability and the developmental stage during which that number is determined. Such work will be relevant to the long-standing question of why some traits display more phenotypic plasticity than others ^133–135^. We suggest that the specific developmental stages during which traits are established may have predictive value in this regard, because different developmental stages interact in distinct ways with the environment.

Among female insects, there is strong evidence that ovariole number correlates positively with reproductive output in some clades, including but not limited to *Drosophila* ^34,136–138,but^ ^see^ ^139^. Ovariole number can also evolve independently of body size ^18,140^, such that insects could in principle evolve higher reproductive outputs without having to evolve larger bodies. Further, when accounting for body size, there is a negative correlation between ovariole number and egg size, which is weak across all studied insect groups ^140^, but very strong in others, including Drosophilidae fruit flies ^136,140^. In some animals, larger eggs can give rise to offspring with higher fitness by metrics including hatching rate ^141^, nutrient contents ^142^ and juvenile size and survival ^143–146^. We previously showed that there is no universal correlation between egg morphology and duration of embryogenesis across all insects ^140^, and studies in some vertebrates and invertebrates show that there is not always a predictable relationship between egg size and offspring fitness metrics ^147–150^. However, this does not preclude the possibility that for specific insect clades, there could be a positive relationship between egg volume and offspring fitness ^151–154^. Although it could seem counter-intuitive to potentially limit offspring number by both reducing ovariole number and eliminating its variance, and thereby its evolvability, we speculate that convergent evolution of invariant gonadiole numbers across insects – rather than simply evolution of lower ovariole numbers ^8^ – might be linked to increased egg size, and in turn, increased offspring fitness. A thorough, phylogenetically informed investigation of the evolutionary and genetic mechanistic relationships between ovariole number, lifetime reproductive capacity, egg size, and offspring fitness across clades showing independent evolution of gonadiole number invariance, would be a valuable avenue for future studies.

## Materials and Methods

### Animal rearing

*O. fasciatus* were maintained as previously described ^21,63,155,156^ on sunflower seeds in the laboratory on a 12h:12h light/dark cycle at 28°C with 70% relative humidity. Since the *Ofas-Blimp1*^22–12^ allele is homozygous lethal, bugs carrying this allele were maintained as a heterozygous stock. Every two or three generations, individual bugs were genotyped using PCR as previously described ^84^ to ensure the mutant allele was maintained. Homozygous mutant embryos were identified in experiments as embryos that had fewer segments and were shorter than wild type embryos; we have independently verified by PCR that all such embryos are indeed homozygous mutant.

### Generation of cladogram with ovariole number in different species of insects

Ovariole and testiole counts from multiple insect species were collated from the literature (Table S1). The evolutionary relationships between the species were drawn using the Interactive Tree of Life v7 ^157^ using the relationships reported by Misof and colleagues ^158^.

### Adult and nymphal dissection and fixation

Adult gonads were dissected from seven-day old adults, a stage by which oogenesis and spermatogenesis have commenced. Gonads were dissected in 1X PBS (phosphate buffered saline) and then imaged in droplets of 1X PBS under white light using a Zeiss SteREO Discovery.V12 or a Zeiss Axio Zoom.V16 stereomicroscope. For immunohistochemistry and *in situ* hybridisation, the sheath surrounding the adult gonads was removed before the gonads were fixed in 4% PFA (paraformaldehyde) in 1X PBS (4% PFA in PBS).

Nymphal instars were identified using characteristic wing morphology as previously described ^159^. Nymphs were pinned, opened along the dorsal midline, eviscerated and fixed in 4% PFA in PBS. Following fixation, gonads were removed from the body cavity for subsequent whole mount immunohistochemistry.

### Embryo collection, fixation and dissection

*O. fasciatus* were allowed to lay eggs in dry cotton for four hours, after which eggs were picked out of the cotton with plastic insect forceps and aged under the same environmental conditions as the animal cultures, until they reached the appropriate stage for analysis. Embryonic staging was based on the developmental series previously described ^63^, expanded and adapted for the present study (Figure S1A; Table S2). All embryonic stages described herein are defined by the range of hours after egg laying (h AEL). Thus, embryos given the same stage number can show all, or only some, of the morphological features described for that stage in Table S2, as they may be up to four hours different in chronological age. *O. fasciatus* eggs of desired ages were stuck to the bottom of a 60 mm Petri dish with double-sided tape and a slit was created in the eggshell with a microscalpel. 4% PFA in PBS was then poured into the dish to fix the embryos for 30 minutes at room temperature, followed by at least three washes with PBT (1X PBS with 0.1% Triton X-100). Embryos were then dissected free of their yolk in PBT with fine forceps. Embryos were washed at least three times with PBT at room temperature before continuing with staining or dehydration in methanol.

### Gene identification

Genes of interest were identified from the *O. fasciatus* genome assembly ^160^ according to existing annotations or based on tBLASTn searches of the genome using homologs from *D. melanogaster* as queries. In all cases, genes for study were considered *O. fasciatus* homologs of the *D. melanogaster* queries, if these sequences were returned as the top hit in reciprocal BLASTp searches. All genes were also amplified from mixed stage embryonic cDNA to verify their sequences using Sanger sequencing (data not shown). When full-length coding sequence was not available, RACE was performed using the SMARTer RACE 5′/3′ Kit (Takara Bio 634859) on mixed stage embryonic cDNA following manufacturer’s instructions. See Table S3.

### Molecular biology

For RNA extraction and cDNA synthesis, embryos or tissues of interest were homogenized in TRIzol (Invitrogen 15596026), and RNA was extracted with phenol–chloroform extraction and isopropanol precipitation. cDNA was synthesized with SuperScript III First-Strand Synthesis SuperMix for qRT-PCR (Invitrogen 11752250) following manufacturer’s instructions.

For cloning and *in vitro* transcription, fragments of genes of interest were PCR-amplified from embryonic cDNA and each cloned into the pGEM-T Easy vector (Promega A137A) using manufacturer’s instructions. A fragment of *GFP* was subcloned from an existing plasmid containing the *GFP* sequence. Insertion was confirmed by Sanger sequencing. Primers used are listed in Table S8.

For *in situ* probe synthesis, gene fragments were PCR-amplified from plasmids using T7 and Sp6 primers. *In vitro* transcription was performed using T7 RNA polymerase (Promega P2075) or Sp6 RNA polymerase (Roche 10810274001) with DIG RNA labelling mix (Roche 11277073910). 100 ng of template was used for a 20-µl reaction run at 37°C overnight. RNA was purified with LiCl/ethanol precipitation, then resuspended in 20 µl DEPC-treated water. Concentration was measured using a NanoDrop Spectrophotometer (Thermo Fisher), and the sample was diluted to 100 ng/µl with hybridisation buffer (50% formamide, 1X PBS, 5x SSC (saline–sodium citrate), 0.1% Triton X-100, 0.1% CHAPS ((3-((3-cholamidopropyl) dimethylammonio)-1-propanesulfonate), 5 mM EDTA (ethylenediaminetetraacetic acid), 50 µg/mL heparin, 1 µg/mL yeast tRNA). Probes were stored at −20°C.

For double-stranded RNA (dsRNA) synthesis, gene fragments were PCR-amplified from plasmids using T7 and T7-Sp6 primers to generate a template for *in vitro* transcription with T7 promoter sequences on both ends. *In vitro* transcription was performed using the MEGAscript T7 Transcription Kit (Invitrogen AM13345). 100 ng of template was used for an 80-µl reaction run at 37°C overnight. RNA was purified with LiCl/ethanol precipitation, then resuspended in 100 µl DEPC (diethyl pyrocarbonate)-treated water. To anneal the two strands, this solution was heated to 95°C on a heat block for 5 minutes, then allowed to cool slowly by leaving the tube on the heat block with the power switched off. A 1:10 dilution of annealed RNA was analysed using a NanoDrop Spectrophotometer (Thermo Fisher) to measure its concentration and purity, then run on a 1% agarose gel to confirm its size and integrity. For RNAi, dsRNA was diluted in DEPC-treated water with 0.05% Texas-Red. Concentrations of injected dsRNA used in each experiment are shown in Tables S5 and S7.

### Search for somatic gonad markers

Somatic gonad precursors in *D. melanogaster* form distinct subpopulations: an anterior population expressing *escargot* (*esg*) and a posterior population expressing *eyes absent* (*eya*) ^9^. *escargot* is one of three *snail*-type transcription factors in *D. melanogaster* ^161,162^^1^^61,162^. Our search of the *O. fasciatus* genome ^160^ yielded only one gene in this family. This gene, *Ofas-snail*, was expressed most highly in the embryonic nervous system (Figure S1C). These data are consistent with previous studies in other arthropods suggesting that the three *D. melanogaster snail-*type family members are the result of gene duplications specific to Diptera ^163–165^. We found that *Ofas-eya* was expressed most strongly in the embryonic optic lobes (Figure S1D). Although both *Ofas*-*snail* and *Ofas-eya* were also expressed in a segmental pattern in the thorax and abdomen in *O. fasciatus*, neither of these expression patterns obviously corresponded to the embryonic gonad, so we did not pursue these candidate genes further.

### Microinjection and RNA interference (RNAi)

*O. fasciatus* were allowed to lay eggs in dry cotton for four hours, after which eggs were picked out of the cotton for injection. Two glass microscope slides were taped together with double-sided tape to create a ledge with exposed double-sided tape against which embryos were lined up. Eggs were then submerged in tap water for at least 20 minutes to soften the eggshell before injection. Slides with injected eggs were kept at 28°C in a humidified dish to continue development until the desired age. Needles for embryonic injection were made by pulling glass capillaries (World Precision Instruments 1B100-4) with a Sutter P-97 Flaming/Brown micropipette puller. Needle tips were manually opened with fine forceps just prior to injection. Eggs were injected with a Narishige IM300 microinjector.

Adult *O. fasciatus* six or more days after the final moult were anesthetised with CO_2_ and injected between the fourth and fifth abdominal segments using a Hamilton syringe and then reared with same-age adults of the opposite sex. Adult RNAi gonad phenotypes were scored five days after injection (Figure S3). Embryonic pRNAi phenotypes were scored in embryos collected within a 24-hour window seven or eight days after injection, and aged until the desired stage of analysis. Concentration of injected dsRNA used in each experiment is shown in Tables S5 and S7.

### Quantitative RT-PCR

Quantitative RT-PCR was performed on an AriaMx Real-time PCR system (Agilent) using PerfeCTa SYBR Green SuperMix, Low ROX (Quantabio 95056). Fold-change differences were calculated using log-transformed relative normalised expression ^166^ for gene expression changes over time. Expression levels were normalised against *Ofas*-*Gapdh1* and *Ofas*-*δ-Tubulin* (See Table S3). All primers used are listed in Table S8.

### Colorimetric in situ hybridisation

Dissected embryos were stored in methanol at −20°C overnight or longer before *in situ* hybridisation. After rehydration in PBT and washing in hybridisation buffer, embryos were hybridised overnight in a 70°C water bath with 1 ng/µl probe in hybridisation buffer. The next day, the following series of 20-minute washes were performed at 70°C: once in hybridisation buffer, three times in wash 1 (2X SSC, 0.1% CHAPS) and three times in wash 2 (0.2X SSC, 0.1% CHAPS). Embryos were then washed three times for ten minutes each at room temperature in KTBT (1X Tris-buffered saline with KCl and 0.1% Triton X-100) followed by an hour-long incubation at room temperature in *in situ* blocking solution (1X PBS, 0.1% Tween-20, 5% normal goat serum, 2 mg/mL bovine serum albumin, 1% dimethyl sulfoxide). Embryos were incubated for two hours at room temperature with sheep anti-digoxigenin–AP (Roche 11093274910), diluted 1:2000 in *in situ* blocking solution, then washed at least four times over two hours with KTBT and incubated in KTBT overnight at 4°C. The following day, embryos were washed with NTMT (100 mM Tris pH 9.5, 100 mM NaCl, 50 mM MgCl_2_, 0.1% Tween-20), then developed in the dark at 37°C with NBT (nitro blue tetrazolium)/BCIP (5-bromo-4-chloro-3-indolyl phosphate) in NTMT with 5% polyvinyl alcohol. Development was monitored at least every hour under a dissection microscope. To stop development of the enzymatic reaction, embryos were washed in PBT, post-fixed in 4% PFA in PBS, then incubated in 70% glycerol before mounting in 100% glycerol.

### Fluorescent hybridisation chain reaction (HCR)

All HCR probes except *Ofas*-*inv* were designed using HCR Probe Maker (Özpolat lab, Washington University St. Louis ^167^) and ordered from IDT as oPools (oligo pools) (Table S9). The *Ofas*-*inv* probes were obtained from Molecular Instruments by submitting a partial cDNA sequence (Table S9). Dissected samples were stored in methanol at −20°C overnight or longer before *in situ* hybridisation. After rehydration in PBT and washing in probe hybridisation buffer (Molecular Instruments), samples were hybridised overnight in a 37°C water bath with 10 nM probe in probe hybridisation buffer. The following day, samples were washed four times for 15 minutes each in probe wash buffer (Molecular Instruments) at 37°C followed by two five-minute washes in SSCT (5X SSC, 0.1% Tween-20) and at least 30 minutes in amplification buffer (Molecular Instruments) at room temperature. Samples were then incubated overnight with 60 nM hairpins (Molecular Instruments) in amplification buffer at room temperature in the dark. On the third day, samples were washed four times for 15 minutes each in SSCT. If applicable, samples were then immunostained as described below. Otherwise, samples were incubated with 2 ng/µl Hoechst 33342 in PBT at room temperature for one hour, post-fixed for 30 minutes in 4% PFA in PBS, then incubated in 70% glycerol before mounting in 100% glycerol.

### Immunohistochemistry

Immunohistochemistry was performed on embryos as previously described ^63^. Primary antibodies used were rabbit anti-Ofas-Vasa (1:100; ^63^), mouse Monoclonal anti-Acetylated Tubulin (6-11B-1, 1:1000; Sigma-Aldrich catalog number T6793) and mouse anti-*Schistocerca gregaria* Pairberry (Pax3/7, DP312, 1:10; ^168^; see also Key Resources Table).

### Imaging and image analysis

Fluorescently labelled tissues were imaged on a Nikon CSU-W1 spinning disc inverted confocal microscope or a Zeiss LSM 980 with Airyscan. Colorimetric *in situ*s were imaged on a Zeiss Axio Observer 7 inverted or a Zeiss Axio Imager Z1 upright compound microscope. Whole mounted adults and dissected and unstained adult gonads were imaged on a Zeiss SteREO Discovery.V12 or a Zeiss Axio Zoom.V16 stereomicroscope. Images were analysed in Fiji v1.54 ^169^ and figures were edited in Inkscape v. 1.1 (retrieved from https://inkscape.org). The SNT Plugin ^170^ in Fiji was used to highlight segments in both the *Ofas*-*gt* RNAi and *Ofas*-*Blimp1* mutant experiments as described below. Zen Blue v. 3.4 (Zeiss) and NIS Elements Viewer v. 5.21.00 (Nikon) were used to visualise data.

### Segment counts in Ofas-gt RNAi and Ofas-Blimp1 experiments

Segment counts were based on *Ofas*-*inv* signal, which marks posterior segment boundaries ^72^. The *Ofas-inv* stripe defining the posterior boundary of T3 was used as the reference point for abdominal segment identification and was identified by the presence of legs anterior to the stripe. All *Ofas-inv* stripes posterior to T3 were considered abdominal and traced toward the posterior end of the embryo. The final analysis focused on the number of segments present within the A1–A7 region, with A7 defined as the most posterior gonadiole-bearing segment. Individual segments were not otherwise assigned specific identities. The total number of segments present was recorded independently for each side of each embryo, along with the corresponding number of gonadiole primordia. Only embryos with clearly visibly thoracic and abdominal morphology and segment boundaries, based on *Ofas-inv* signal, were used for analyses.

## Key Resources Table

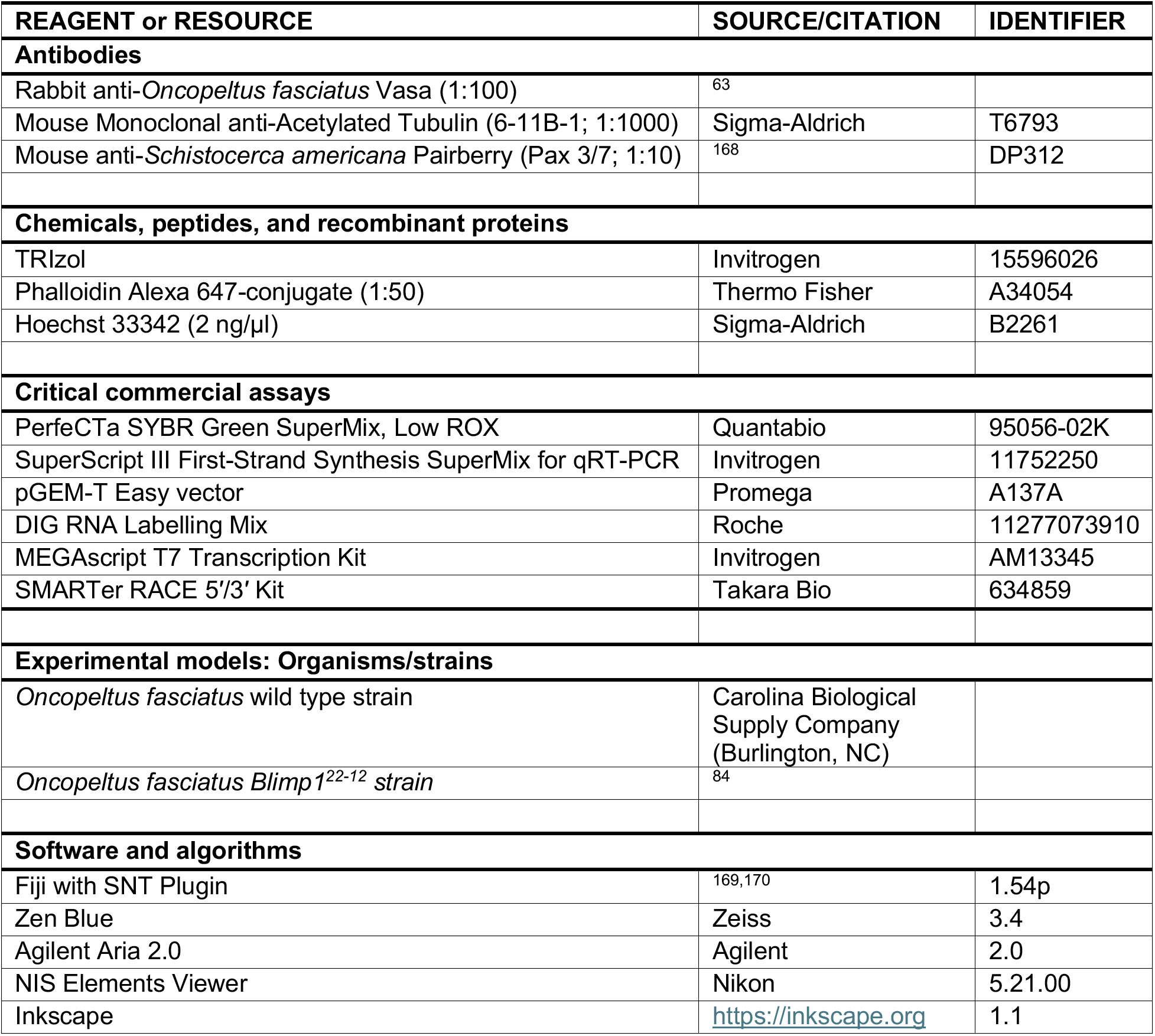

## Acknowledgements

The authors thank Extavour lab members for helpful discussions, Katie Reding and Leslie Pick (University of Maryland, College Park, MD, USA) for the *Ofas-Blimp1*^22–12^ mutant line, and Nipam Patel (Marine Biological Laboratory, Woods Hole, MA, USA) for the anti-Pax3/7 (DP312) antibody. This work was supported by National Science Foundation award IOS-2220747, funds from Harvard University, and funds from the Howard Hughes Medical Institute (HHMI) to CGE. CGE is an HHMI Investigator.

## Author Contributions

TK and CGE conceived of the study. JAK, TK and DAH performed experiments. All authors analysed data. JAK and TK wrote the first draft of the manuscript. All authors reviewed and edited the manuscript. CGE obtained funding for and oversaw the study.

## Generative AI Use Statement

Chat GPT (5.6 Sol Light) was used occasionally as an initial literature search engine to find scientific papers, which were then manually verified for veracity by the authors. It was also used as an interface to ask about clarity of text that had been manually written by the authors. No generative AI was used to generate content of any kind.

## Supplemental Information

**Figure S1.**
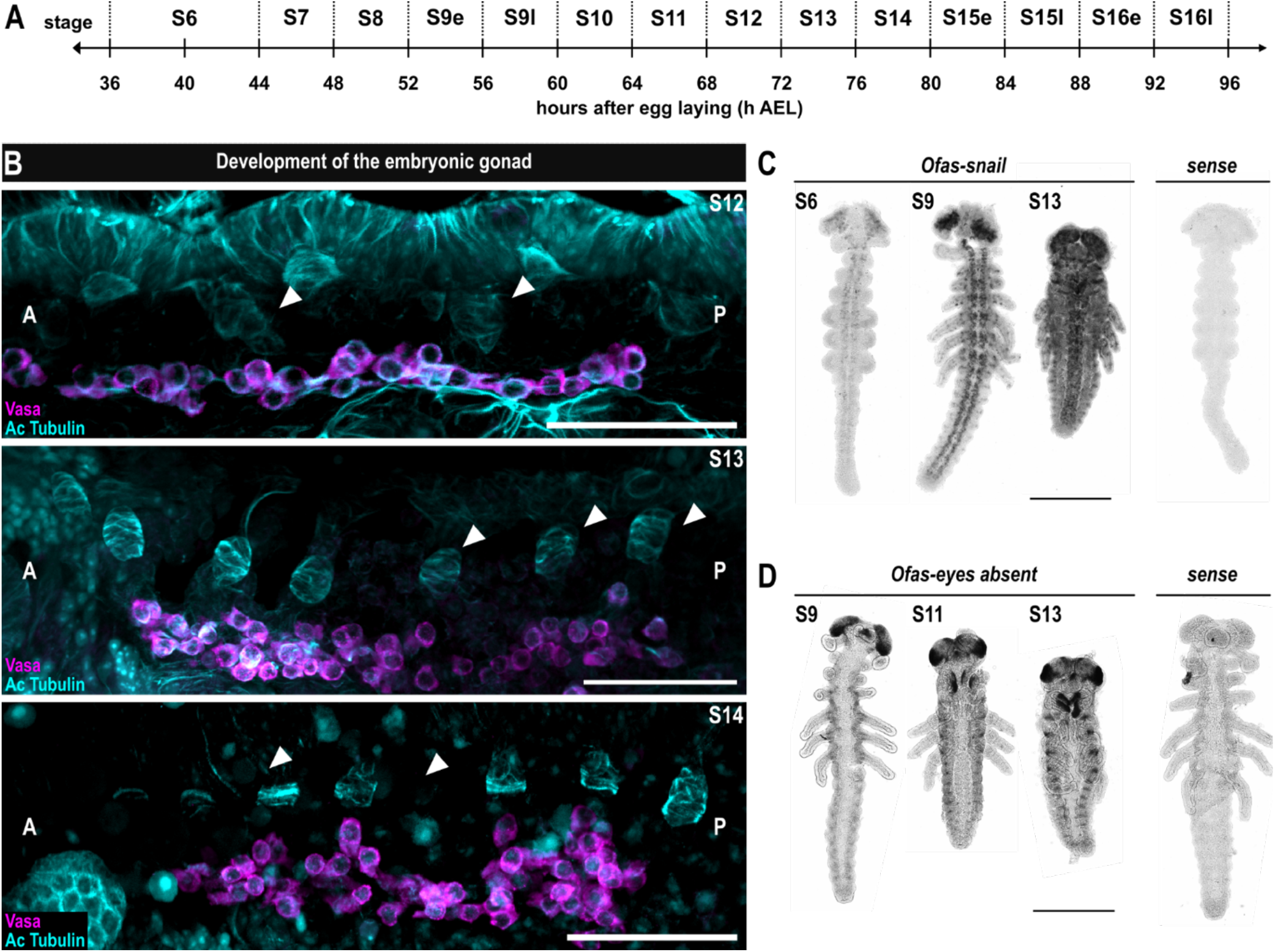
Acetylated Tubulin is expressed strongly in the presumptive embryonic somatic gonad primordia and weakly in the germ cells. (**A**) Embryonic stages used in this study, indicating ranges of developmental time (hours after egg laying (h AEL) spanned by each stage. (See also Table S2). (**B**) Cells in seven lateral abdominal domains expressing acetylated Tubulin (Ac Tubulin, cyan) are shown associated with the germ cells expressing Ofas-Vas (magenta) across three sequential stages of embryonic development. White arrowheads point to individual gonad precursors. A and P depict the anteroposterior axis of the embryo. Scale bars = 50 µm. (**C-D**) Colorimetric *in situ* hybridisation of (C) *Ofas*-*snail* (*snail*) and (D) *Ofas*-*eyes absent* (*eya*) across three stages of embryonic development. Scale bars = 1000 µm.

**Figure S2.**
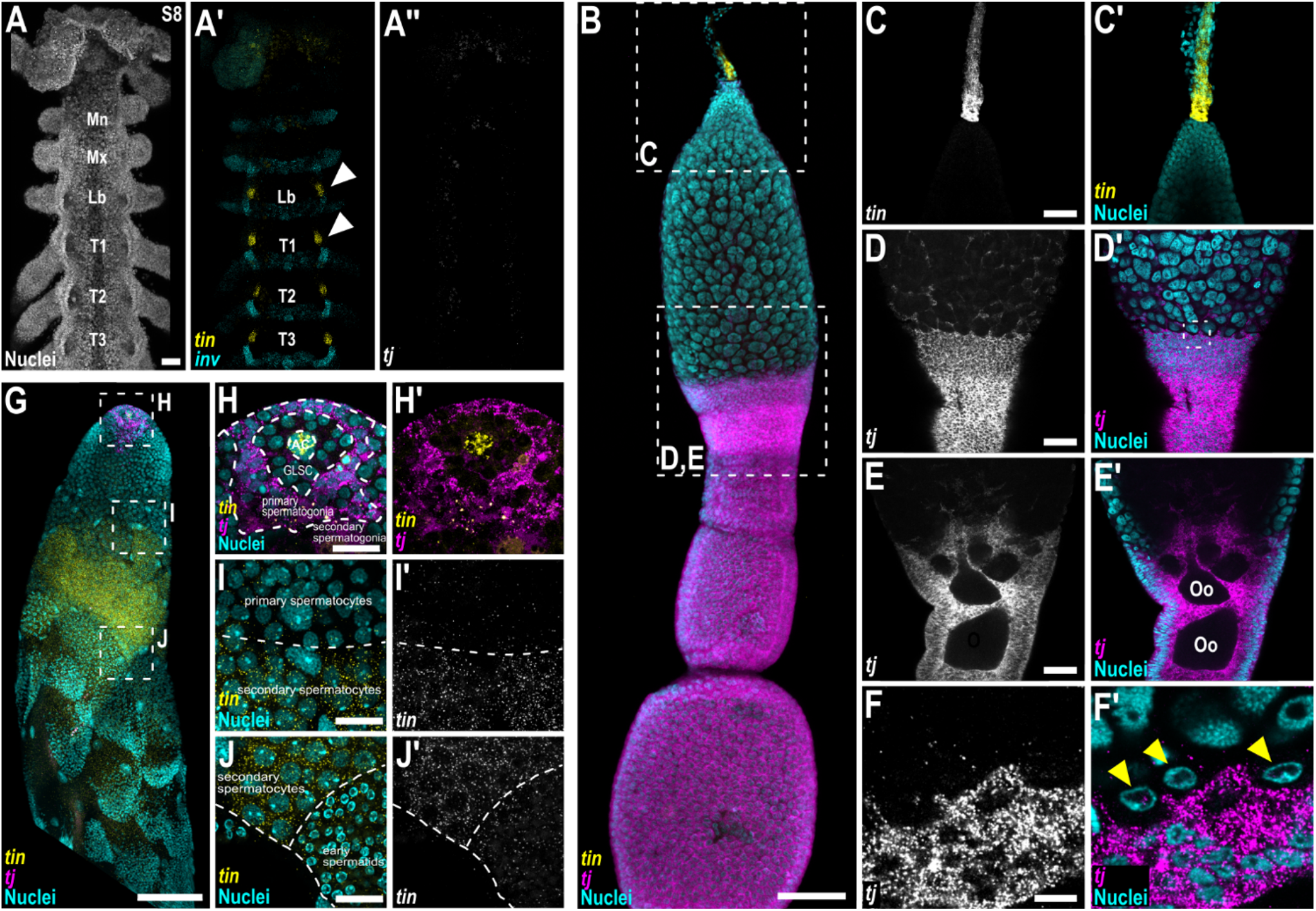
*Ofas-tj* and *Ofas-tin* are expressed in the adult gonads. **(A**) Head and thoracic segments of a stage 8 embryo. (**A’**) *Ofas*-*tin* (yellow) marks heart precursors in all segments including and posterior to the labial segment. *Ofas*-*inv* (cyan) marks the posterior compartment of each segment. (**A’’**) *Ofas*-*tj* (magenta) expression is not detected in the head or thoracic segments. Abdominal expression of *Ofas*-*tj* and *Ofas*-*tin* at this stage is shown in Figure 2A. Nuclei stained with Hoechst 33342 (grey) show embryonic morphology. Segment identity is indicated in each panel. Scale bar = 50 µm. **(B)** In adult ovaries, *Ofas*-*tj* was expressed in follicle cells surrounding developing oocytes throughout oogenesis, whereas *Ofas*-*tin* expression was detected in the terminal filament of each ovariole. Scale bar = 100 µm. **(C-E)** Higher magnification of boxed regions in (B) showing *Ofas*-*tin* and *Ofas*-*tj* expression in distinct regions of the ovariole: *Ofas*-*tin* is expressed in the terminal filament (**C-C’**). *Ofas*-*tj* is expressed in follicle cells of both the tropharium (**D-D’**) and developing oocytes (**E-E’**; Oo). Scale bars = 25 μm. *Ofas*-*tj* expression is not detected in late-stage oocytes (E-E’) or early oocytes (**F-F’**; yellow arrowheads). Scale bar = 50 μm. **(G)** In adult testes, *Ofas*-*tin* was expressed in the apical complex of each testiole, whereas *Ofas*-*tj* was detected in the somatic cells surrounding the primary spermatogonia. In addition to its apical expression, *Ofas*-*tin* was strongly expressed in a discrete region near the middle of the testiole corresponding to the secondary spermatocytes. Scale bar = 100 µm. **(H-J)** Higher magnification of insets in (G) showing *Ofas*-*tin* and *Ofas*-*tj* expression in distinct regions of the testioles. *Ofas*-*tj* is expressed in the region surrounding the germline stem cells (GLSC) where both presumptive gonialblasts and cyst progenitor cells can be found, whereas *Ofas*-*tin* is expressed in the apical complex and in secondary spermatocytes. The stages of spermatogenesis ^171,172^ are labelled. Scale bars = 25 μm.

**Figure S3.**
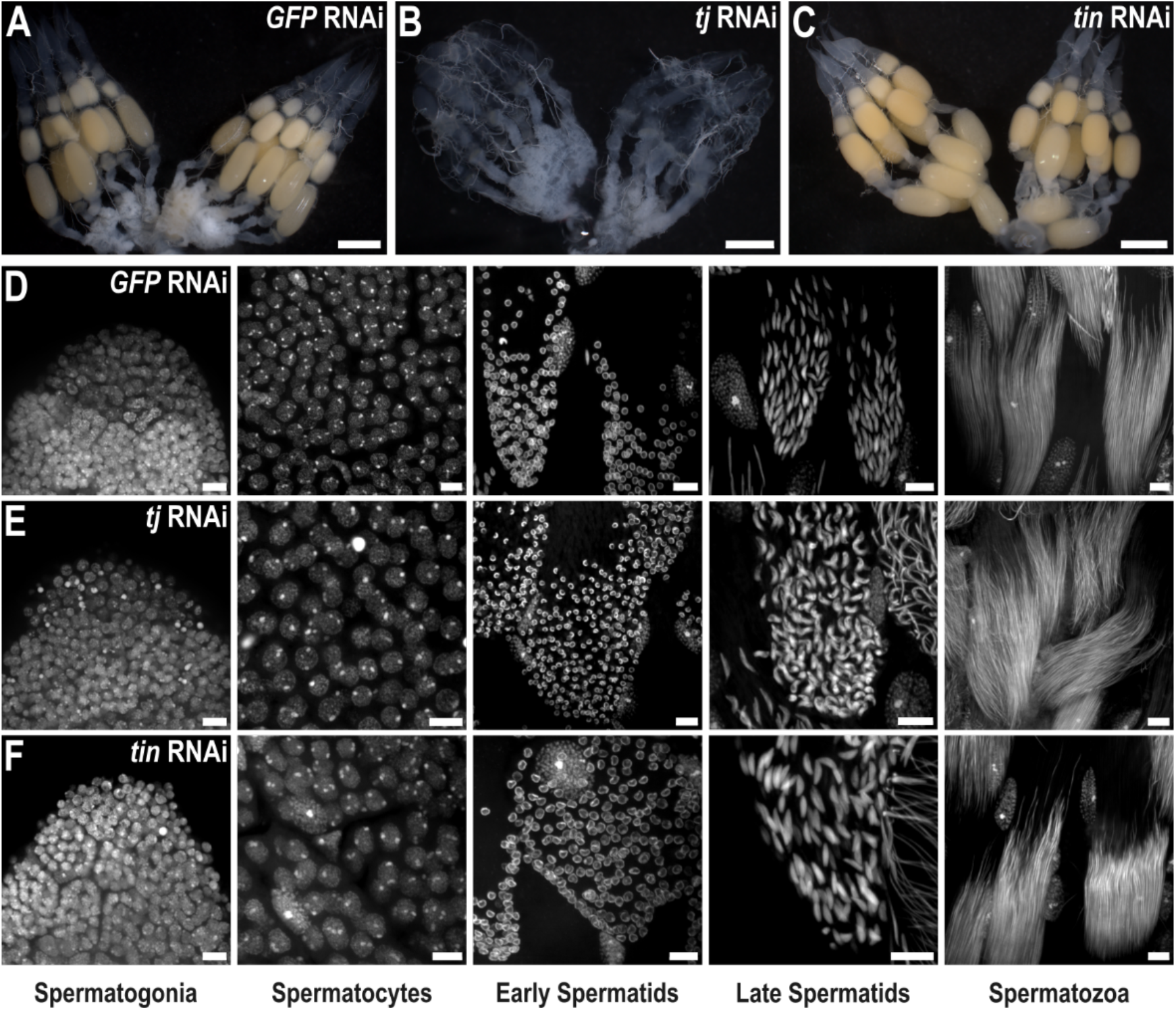
*Ofas*-*tj* is required for oogenesis. We performed RNAi against *Ofas*-*tin* and *Ofas*-*tj* in adult females and males. Reduction of *Ofas*-*tin* produced no readily detectable defects in oogenesis (A, C), whereas *Ofas*-*tj* RNAi resulted in a failure of oogenesis (B), suggesting a requirement for *Ofas*-*tj* during female gametogenesis under these conditions. **(A-C)** Representative ovaries, five days after injection, from adult females injected with (A) *GFP* dsRNA (*GFP* RNAi, n=12), (B) *Ofas*-*tj* dsRNA (*tj* RNAi, n=16) or (C) *Ofas*-*tin* dsRNA (*tin* RNAi, n=18). Scale bars = 1000 µm. By contrast, neither *Ofas*-*tin* nor *Ofas*-*tj* RNAi produced an overt defect in spermatogenesis: all morphological stages of spermatogenesis remained detectable in dsRNA-injected males. **(D-F)** Representative testes from adult males injected with (D) dsRNA targeting *GFP* (*GFP* RNAi, n=24), (E) *Ofas*-*tj* (*tj* RNAi, n=37), or (F) *Ofas*-*tin* (*tin* RNAi, n=29). Testes were stained with Hoechst 33342 (grey) to visualise the stages of spermatogenesis, which appear unaffected by either *Ofas*-*tj* or *Ofas*-*tin* RNAi. Scale bars = 10 μm.

**Figure S4.**
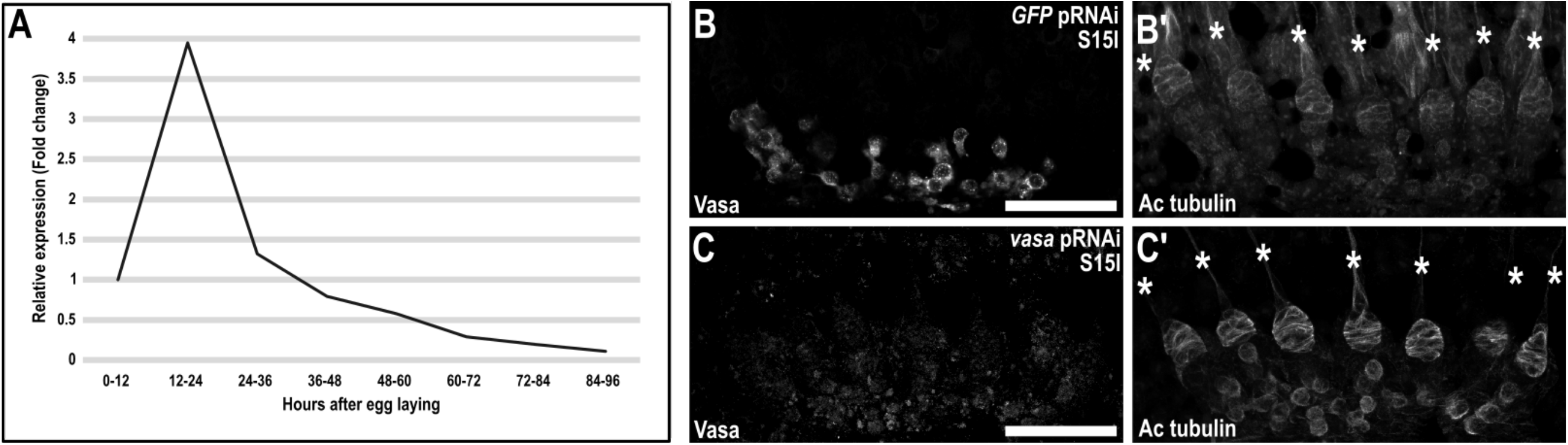
Maternal *Ofas*-*vas* RNAi reduces *Ofas*-*vas* expression but does not prevent formation of gonadiole primordia. **(A)** Relative *Ofas*-*vas* expression levels across sequential stages of wild type embryonic development quantified by qPCR and presented as fold change relative to embryos collected 0-12 hours after egg laying. **(B-C’)** Representative gonads in stage 15 embryos laid by mothers injected with (B-B’) *GFP* dsRNA (*GFP* pRNAi) or (C-C’) *Ofas*-*vas* dsRNA (*vasa* pRNAi), stained with anti-Ofas-Vasa (grey) to detect Ofas-Vas protein levels, Acetylated Tubulin (Ac Tubulin, green) to highlight somatic gonadal structures and germ cells. Asterisks indicate individual gonadiole primordia. Scale bars = 50 µm.

**Figure S5.**
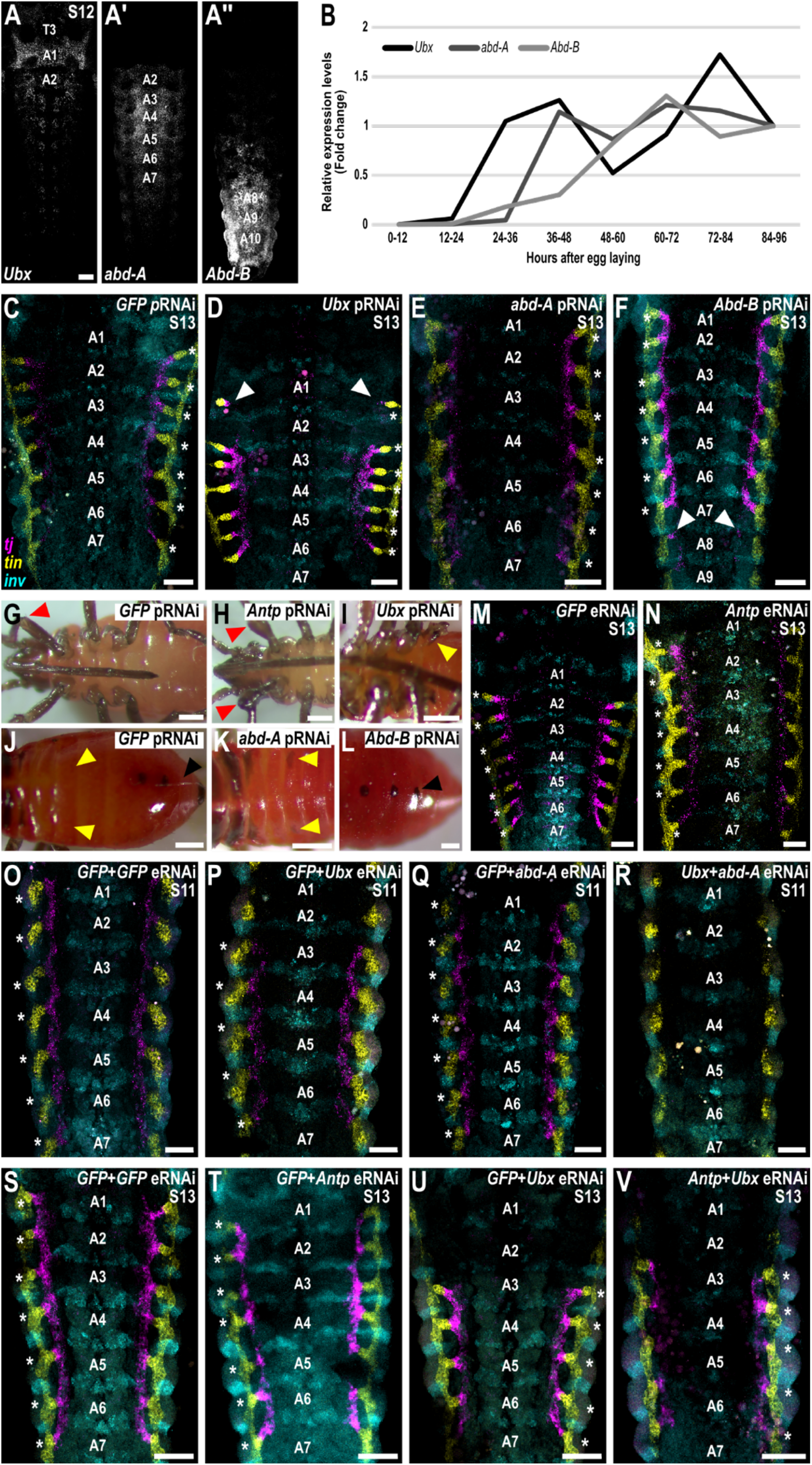
Posterior Hox genes regulate gonadiole primordia formation (previous page). **(A)** Expression of *Ofas*-*Ultrabithorax* (*Ubx*), *Ofas*-*abdominal A* (*abd-A*) and *Ofas*-*Abdominal B* (*Abd-B*) in the stage 12 embryo shown in Figure 5A, displayed as individual channels. **(B)** Relative *Ofas*-*Ubx, Ofas*-*abd-A* and *Ofas*-*Abd-B* expression levels across sequential stages of embryonic development, quantified by qPCR and presented as fold change relative to expression in eggs collected 84-96 hours after egg laying. **(C-F)** Representative images of selected abdominal segments of stage 13 embryos from adult females injected with (C) dsRNA targeting *GFP* (*GFP* pRNAi), (D) *Ofas*-*Ubx* (*Ubx* pRNAi), (E) *Ofas*-*abd-A* (*abd-A* pRNAi), or (F) *Ofas*-*Abd-B* (*Abd-B* pRNAi). In *Ofas*-*Ubx* pRNAi embryos, the A1 gonadiole appears to form (white arrowheads) but does not connect to the rest of the gonad (n=11/25). In *Ofas*-*Abd-B* pRNAi embryos, *Ofas*-*tj* is expressed ectopically in A8 (red arrowheads, n=5/16). *GFP* eRNAi (n=42/42) and *Ofas*-*abd-A* pRNAi (n=30/30) embryos show no change in somatic gonad precursor specification. Asterisks indicate individual gonadiole primordia. Scale bars = 50 µm. **(G-I)** Segment transformations in nymphs hatched from eggs laid by adult females injected with (G) dsRNA targeting *GFP* (*GFP* pRNAi, n=30/30), (H) *Ofas*-*Antp* (*Antp* pRNAi; n=5/11) and (I) *Ofas*-*Ubx* (*Ubx* pRNAi; n=4/11). Ventral views are shown with the anterior to the left. Red arrowheads indicate the distal femur of the first leg, and yellow arrowheads indicate ectopic leg-like appendages. Scale bars = 200 μm. **(J-L)** Segment transformation in nymphs hatched from eggs laid by adult females injected with *dsRNA* targeting (J) *GFP* (*GFP* pRNAi, n=30/30), (K) *Ofas*-*abd-A* (*abd-A* pRNAi; n=3/11) and (L) *Ofas*-*Abd-B* (*Abd-B* pRNAi; n=2/20). Dorsal views of the abdomen are shown with the anterior to the left. Yellow arrowheads indicate abdominal segment A3, and black arrowheads indicate abdominal segment A7. Scale bars = 200 μm. **(M-N)** Representative images of selected abdominal segments of stage 13 embryos injected with dsRNA targeting (M) *GFP* (*GFP* eRNAi, n=69/69) and (N) *Ofas*-*Antp* (*Antp* eRNAi; n=35/35). There is no apparent change in gonadiole formation in either condition. **(O-R)** Representative images of selected abdominal segments of stage 11 embryos injected with dsRNA targeting (O) *GFP+ GFP* double eRNAi (*GFP*+*GFP* eRNAi), (P) *GFP* + *Ofas*-*Ubx* (2 µg/µl each), (Q) *GFP*+ *Ofas*-*abd-A* (2 µg/µl each) and (R) *Ofas*-*Ubx*+ *Ofas*-*abd-A* (2 µg/µl each). **(S-V)** Representative images of selected abdominal segments of stage 13 embryos injected with dsRNA targeting (S) *GFP+ GFP* double eRNAi (*GFP*+*GFP* eRNAi), (T) *GFP*+ *Ofas*-*Antp* (2 µg/µl each), (U) *GFP+Ofas*-*Ubx* (2 µg/µl each) and (V) *Ofas*-*Antp*+ *Ofas*-*Ubx* (2 µg/µl each). Additional knockdown against *Ofas*-*Antp* does not change the *Ofas*-*Ubx* knockdown phenotype. In panels **(C-F; M-V)**, *Ofas*-*tj* (magenta) and *Ofas*-*tin* (*y*ellow) mark somatic gonad precursors. *Ofas*-*inv* (cyan) marks the posterior compartment of each segment. Scale bars = 50 µm.

**Figure S6.**
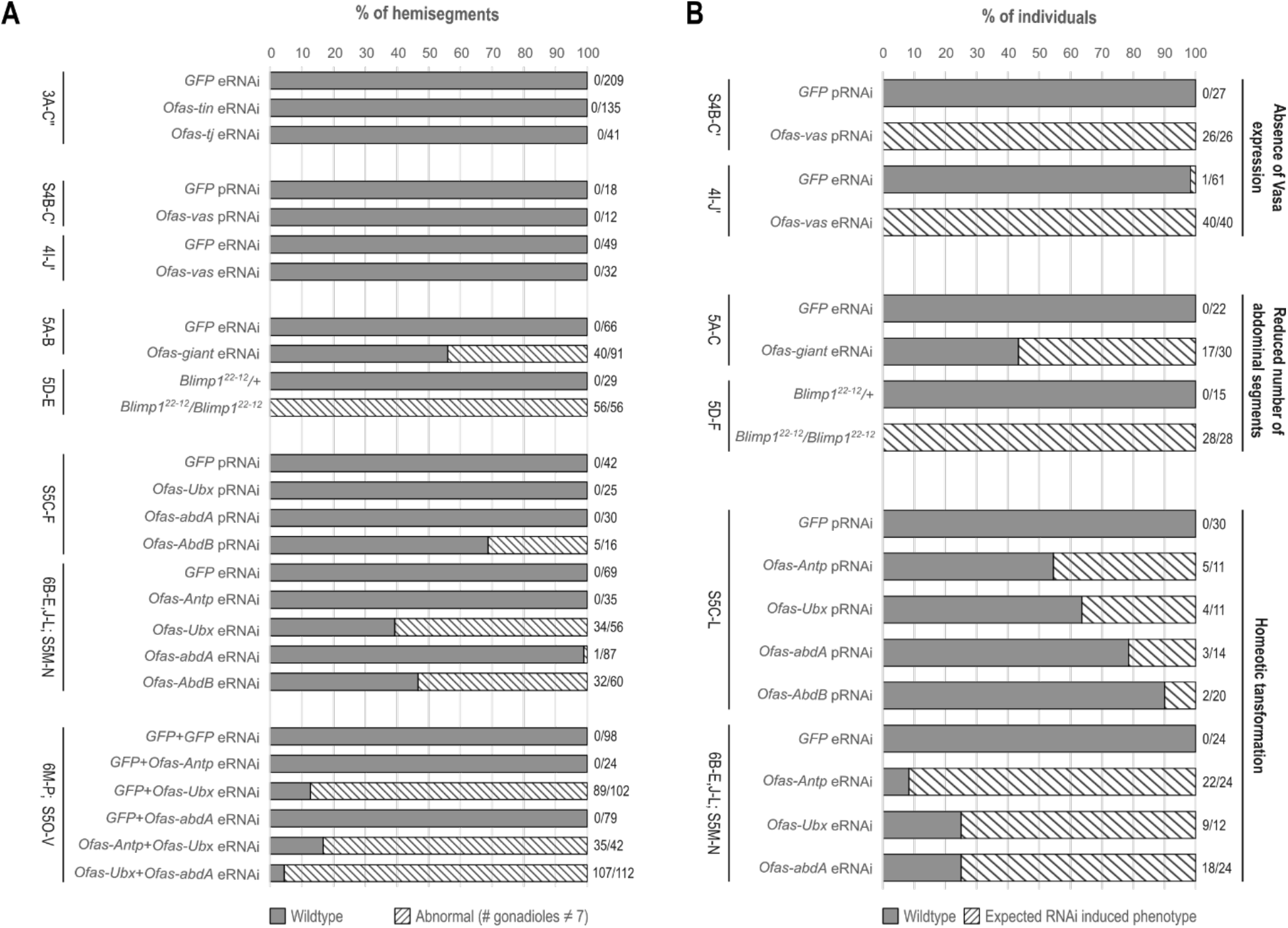
Phenotypic penetrance of dsRNA-induced phenotypes. **(A)** Bar plots showing the percentage of hemisegments containing an abnormal number of gonadioles (n ≠ 7) for each experimental manipulation. The corresponding figure number is indicated to the left of each plot, and the ratio of abnormal hemisegments to the total number of hemisegments analysed is indicated to the right. Abnormal hemisegments are shown as a hatched bar while wild type hemisegments are shown as a solid grey bar. **(B)** Bar plots showing the penetrance of expected dsRNA induced phenotypes as a percentage of the total number of individuals examined. The efficacy of dsRNA was assessed as the percentage of embryos lacking detectable Ofas-Vas protein expression in the germ cells (absence of Ofas-Vas expression; Figure 4I-J’, Figure S4B-C’). The efficacy of *Ofas-gt* dsRNA and the penetrance of the *Ofas-Blimp1*^22–12^ homozygous phenotype were assessed as the percentage of embryos with a reduced number of abdominal segments (Reduced number of abdominal segments; Figure 5A-D). The efficacy of Hox gene dsRNA was assessed as the percentage of embryos exhibiting homeotic transformations, as described in Table S6. Efficacy was not quantified following embryonic injection of *Ofas-Abd-B* dsRNA because the injected embryos did not hatch and lacked an easily identifiable embryonic phenotype. The corresponding figure number is indicated to the left of each plot, and the ratio of individuals with expected dsRNA phenotypes to the total number of individuals analysed is indicated to the right. dsRNA-induced phenotypes are shown as a hatched bar; wild type individuals are shown as a solid grey bar.

**Table S1.** References for gonadial numbers presented in. **Figure 1A**. This list includes the literature sources from which ovariole and testiole numbers were obtained.

| **Species** | **Ovariole references** | **Testiole references** |
| --- | --- | --- |
| *Thermobia domestica* | 57,173–175 | 57,173,176 |
| *Cloeon dipterum* | 177–179 | 180 |
| *Gryllus campestris* | 181 | 182 |
| *Gryllus assimilis* | 183 | 183 |
| *Periplaneta americana* | 184 | 185 |
| *Oncopeltus fasciatus* | 60,61 | 60,61 |
| *Nasonia vitripennis* | 186 | 187 |
| *Tribolium castaneum* | 188–191 | 192–195 |
| *Bombyx mori* | 196,197 | 196,198 |
| *Anopheles gambiae* | 199 | 200 |
| *Aedes aegypti* | 201,202 | 203,204 |
| *Drosophila sechellia* | 19,31,38,39,205,206 | 207 |
| *Drosophila melanogaster* | 19,31,38 | 207,208 |
| *Drosophila virilis* | 209 | 210 |

**Table S2.**
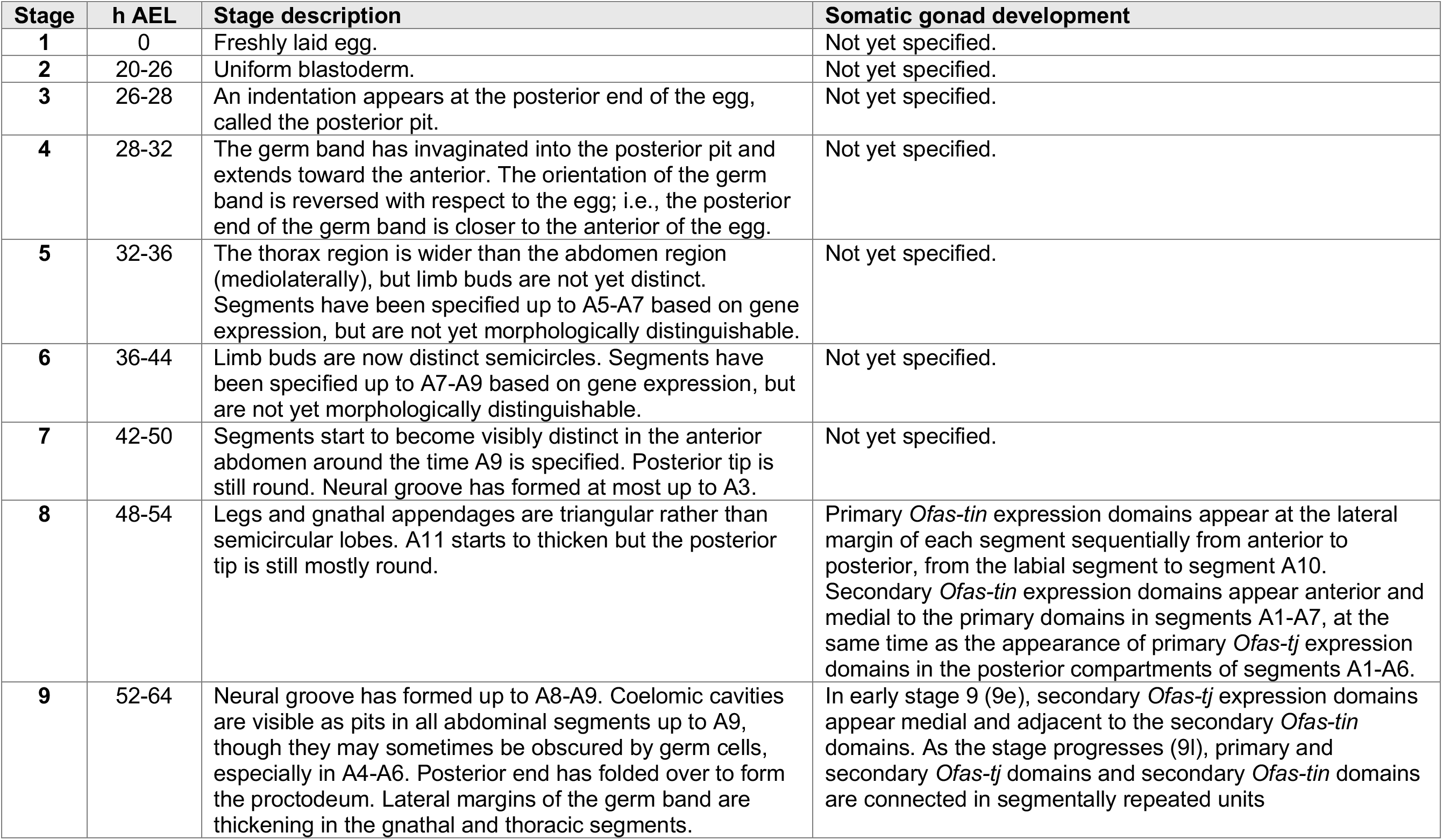
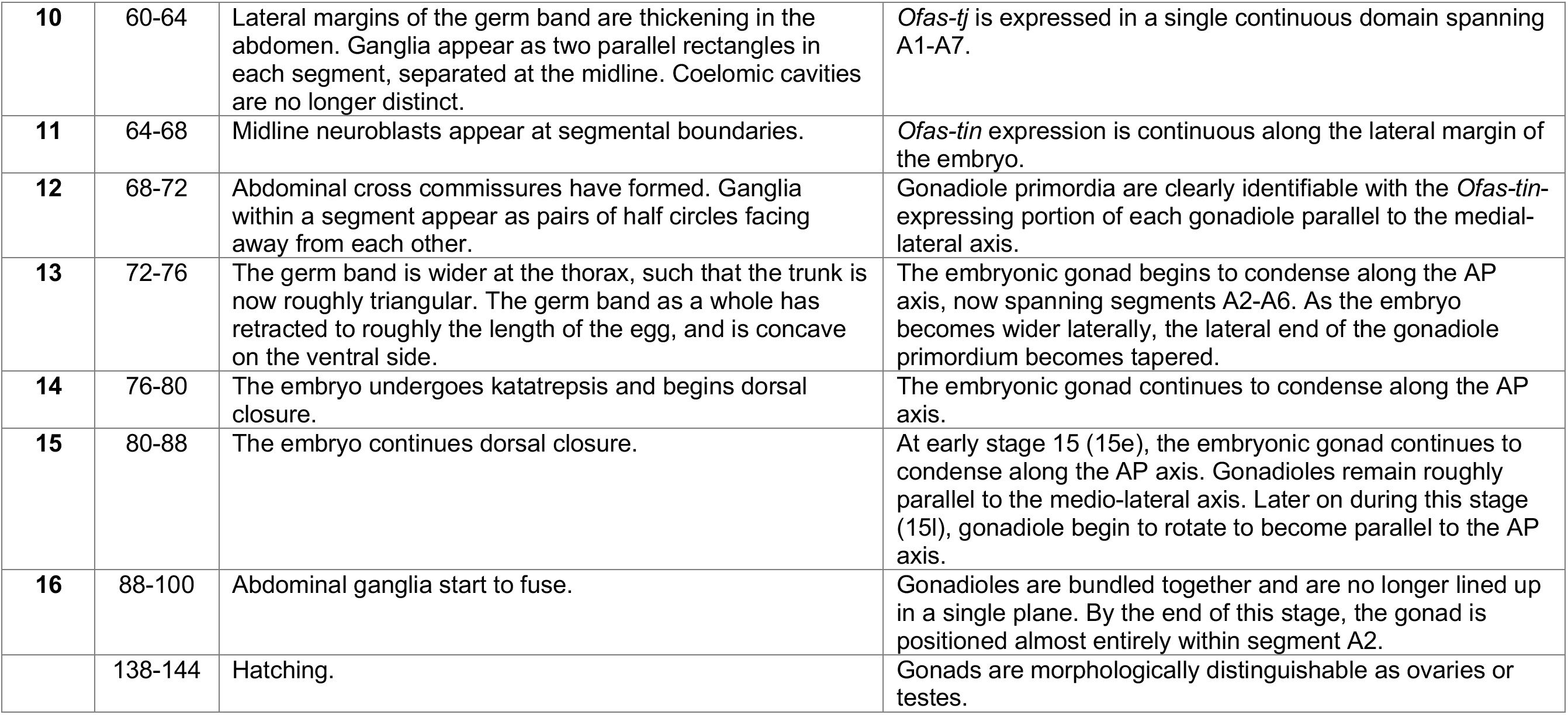
Staging table for stages of wild type *O. fasciatus* embryonic development considered in this study. Descriptions of embryonic stages, adapted from ^63^ with corresponding descriptions of somatic gonad development, through to 100 hours after egg laying (h AEL). All times correspond to wild type development at 28°C and 70% relative humidity.

**Table S3.** List of analysed genes. Genes analysed in this study, identified by their gene ID and scaffold number in Official Gene Set 1.2 (accessed between 10 March 2022 and 14 September 2024) of the *O. fasciatus* genome ^160^. The “Annotated?” column indicates whether the gene was already previously identified and named in Official Gene Set 1.2. The “RACE?” column indicates whether RACE was performed to obtain full-length coding sequence. Full-length sequences obtained through RACE were deposited to GenBank and can be accessed with the accession numbers listed in the “GenBank Accession” column.

| *O. fasciatus* gene name | Gene ID | Scaffold # | Annotated? | RACE? | GenBank accession |
| --- | --- | --- | --- | --- | --- |
| *Ofas-snail (sna)* | OFAS025184 | 1781 | yes | yes | PZ910024 |
| *Ofas-eyes absent (eya)* | OFAS018834 | 4793 | yes | yes | PZ910023 |
| *Ofas-tinman (tin)* | OFAS005760 | 212 | yes | yes | PZ276395 |
| *Ofas-traffic jam (tj)* | OFAS000984 | 14 | no | yes | PZ276396 |
| *Ofas-invected (inv)* | OFAS025056 | 112 | yes | no |  |
| *Ofas-giant (gt)* | OFAS025035 | 3738 | yes | no |  |
| *Ofas-Antennapedia (Antp)* | OFAS006718 | 307 | yes | no |  |
| *Ofas-Ultrabithorax (Ubx)<sup>a</sup>* | OFAS025026 | 3423 | yes | no |  |
|  | OFAS004892 | 220 |  |  |  |
| *Ofas-abdominal A (abd-A)* | OFAS025133 | 1365 | yes | no |  |
| *Ofas-Abdominal B (Abd-B)* | OFAS025323 | 293 | yes | no |  |
| *Ofas-Gapdh1* | OFAS009253 | 1053 | no | no |  |
| *Ofas-δ-Tubulin<sup>b</sup>* | OFAS012604 | 982 | no | no |  |

<sup>a</sup> The *Ofas-Ubx* gene model is split across two different scaffolds which have not yet been assembled together as of the most recent release. 025026 is exon 1 (5' end) and 004892 is exon 2 (3' end). The dsRNA sequence used in this study falls entirely within exon 1.
<sup>b</sup> The *Ofas-d-Tubulin* gene we report here, which was used for qPCR normalization (see Table S8), was amplified with primers that we previously<sup>79</sup> reported as amplifying *Ofas- $\beta$ -Tubulin*, before the current genome assembly<sup>160</sup> was available. Based on the current genome assembly, we now believe that the primers used in this study amplify *Ofas-d-Tubulin*, as all reciprocal BLAST hits of the amplicon obtained with these primers are identified as $\delta$ -tubulin.

**Table S4.**
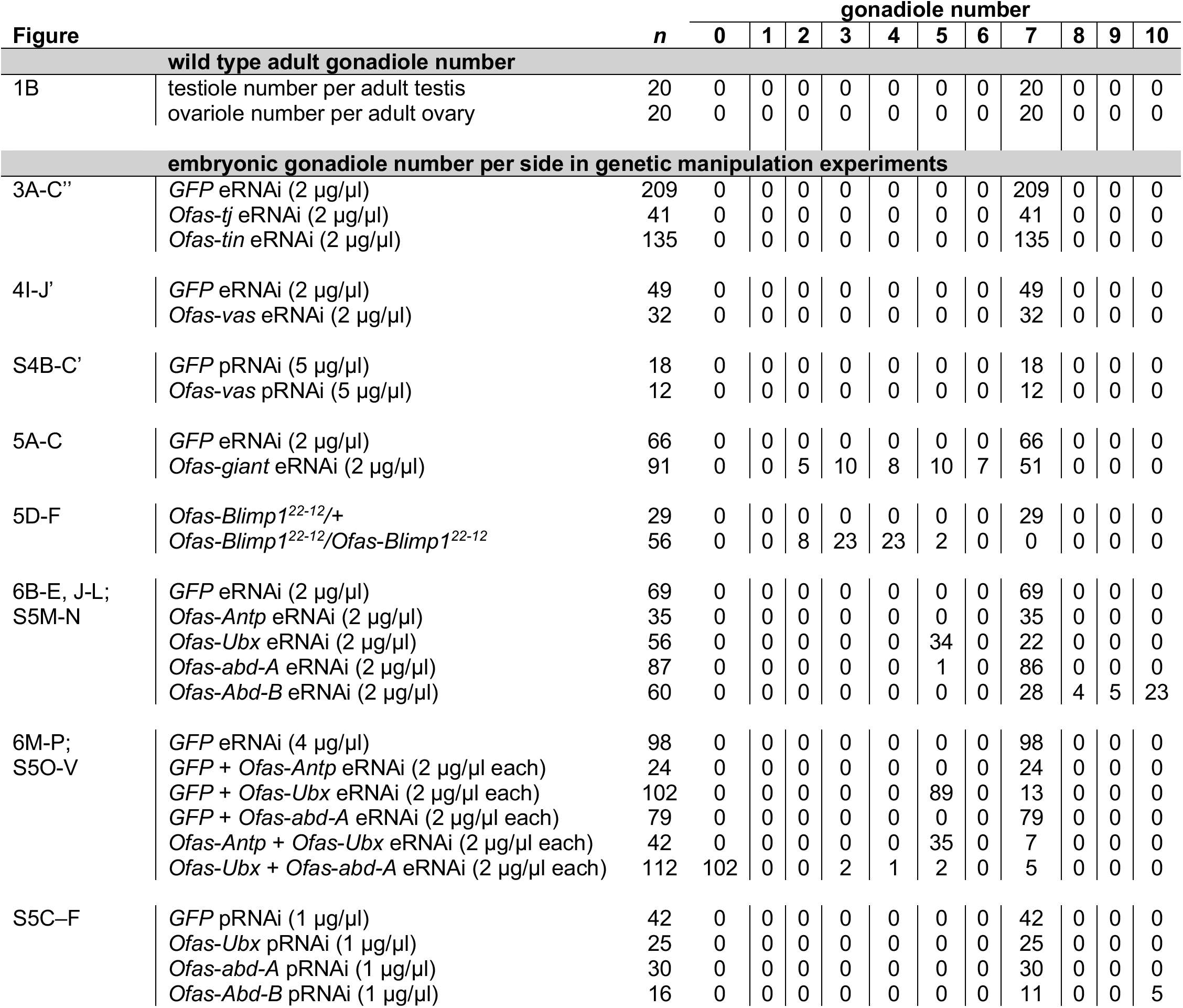
Raw gonadiole counts. Gonadiole counts across various experimental conditions described in this study. For embryonic gonadiole number quantification, each data point represents the number of gonadioles on either the left or the ride side of an embryo (corresponding to the number of gonadioles that would be in one adult gonad).

**Table S5.** Embryonic injection statistics. Survival statistics for embryonic injection experiments. All embryos were injected at 0-4 hours after egg laying. An embryo was scored as “survived” if it was still intact 24 hours after injection.

| Figure | dsRNA | Concentration (µg/µl) | # injected | # survived | % survival |
| --- | --- | --- | --- | --- | --- |
| 3A-C'' | *GFP* | 2 | 444 | 357 | 80.41 |
|  | *Ofas-tj* | 2 | 114 | 67 | 58.77 |
|  | *Ofas-tin* | 2 | 470 | 388 | 82.55 |
| 3D-G'' | *GFP* | 4 | 106 | 86 | 81.13 |
|  | *GFP* + *Ofas-tin* | 2 each | 105 | 104 | 99.05 |
|  | *GFP* + *Ofas-tj* | 2 each | 104 | 103 | 99.04 |
|  | *Ofas-tin* + *Ofas-tj* | 2 each | 159 | 147 | 92.45 |
| 4I-J' | *GFP* | 2 | 119 | 117 | 98.32 |
|  | *Ofas-vas* | 2 | 114 | 89 | 78.07 |
| 5A-C | *GFP* | 2 | 392 | 330 | 84.18 |
|  | *Ofas-gt* | 2 | 510 | 465 | 91.18 |
| 6B-E, J-L;<br>S5M-N | *GFP* | 2 | 615 | 558 | 90.73 |
|  | *Ofas-Antp* | 2 | 113 | 109 | 96.46 |
|  | *Ofas-Ubx* | 2 | 450 | 397 | 88.22 |
|  | *Ofas-abd-A* | 2 | 559 | 485 | 86.76 |
|  | *Ofas-Abd-B* | 2 | 437 | 406 | 92.91 |
| 6M-P;<br>S5O-V | *GFP* | 4 | 355 | 306 | 86.20 |
|  | *GFP* + *Ofas-Antp* | 2 each | 123 | 120 | 97.56 |
|  | *GFP* + *Ofas-Ubx* | 2 each | 355 | 326 | 91.83 |
|  | *GFP* + *Ofas-abd-A* | 2 each | 353 | 344 | 97.45 |
|  | *Ofas-Antp* + *Ofas-Ubx* | 2 each | 128 | 110 | 85.94 |
|  | *Ofas-Ubx* + *Ofas-abd-A* | 2 each | 354 | 337 | 95.20 |

**Table S6.** Penetrance of dsRNA-induced phenotypes. Phenotypes scored in addition to gonadiole primordia number in embryos and hatchlings following dsRNA treatments. For Hox gene knockdown experiments, representative images of homeotic transformations are included in Figure S5G–H”, and are consistent with those described in previous reports ^86^.

| Figure | Condition | # embryos analysed | # displaying phenotype | % penetrance | Observed phenotype |
| --- | --- | --- | --- | --- | --- |
| 4I-J' | *GFP* eRNAi (2 µg/µl) | 61 | 1 | 1.64 | NA |
|  | *Ofas-vas* eRNAi (2 µg/µl) | 40 | 40 | 100.00 | Absence of Vasa expression |
| S4B-C' | *GFP* pRNAi (5 µg/µl) | 27 | 0 | 0.00 | NA |
|  | *Ofas-vas* pRNAi (5 µg/µl) | 26 | 26 | 100.00 | Absence of Vasa expression in laid embryos |
| 5A-C | *GFP* eRNAi (2 µg/µl) | 22 | 0 | 0.00 | NA |
|  | *Ofas-gt* eRNAi (2 µg/µl) | 30 | 17 | 56.67 | Reduction in the total number of abdominal segments |
| 5D-F | *Ofas-Blimp1*<sup>22-12/+</sup> | 15 | 0 | 0.00 | NA |
|  | *Ofas-Blimp1*<sup>22-12</sup>/*Ofas-Blimp1*<sup>22-12</sup> | 28 | 28 | 100.00 | Reduction in the total number of abdominal segments |
| 6B-E,J-L; | *GFP* eRNAi (2 µg/µl) | 24 | 0 | 0.00 | NA |
| S5M-N | *Ofas-Antp* eRNAi (2 µg/µl) | 24 | 22 | 91.67 | Complete transformation of all thoracic limbs to antennae |
|  | *Ofas-Ubx* eRNAi (2 µg/µl) | 12 | 9 | 75.00 | Legs on segment A1 |
|  | *Ofas-abd-A* eRNAi (2 µg/µl) | 24 | 18 | 75.00 | Ectopic appendages on segments A2–A8 |
|  | *Ofas-Abd-B* eRNAi (2 µg/µl) | NA | NA | NA | No visible embryonic phenotype prior to hatching; embryos do not hatch |
| S5C-L | *GFP* pRNAi (1 µg/µl) | 30 | 0 | 0.00 | NA |
|  | *Ofas-Antp* pRNAi (1 µg/µl) | 11 | 5 | 45.45 | Swelling of the distal femur |
|  | *Ofas-Ubx* pRNAi (1 µg/µl) | 11 | 4 | 36.36 | Partial legs on segment A1 |
|  | *Ofas-abd-A* pRNAi (1 µg/µl) | 14 | 3 | 21.43 | Pigmentation in A5/A6 |
|  | *Ofas-Abd-B* pRNAi (1 µg/µl) | 20 | 2 | 10.00 | Ectopic pigmentation on segment A7 |

**Table S7.**
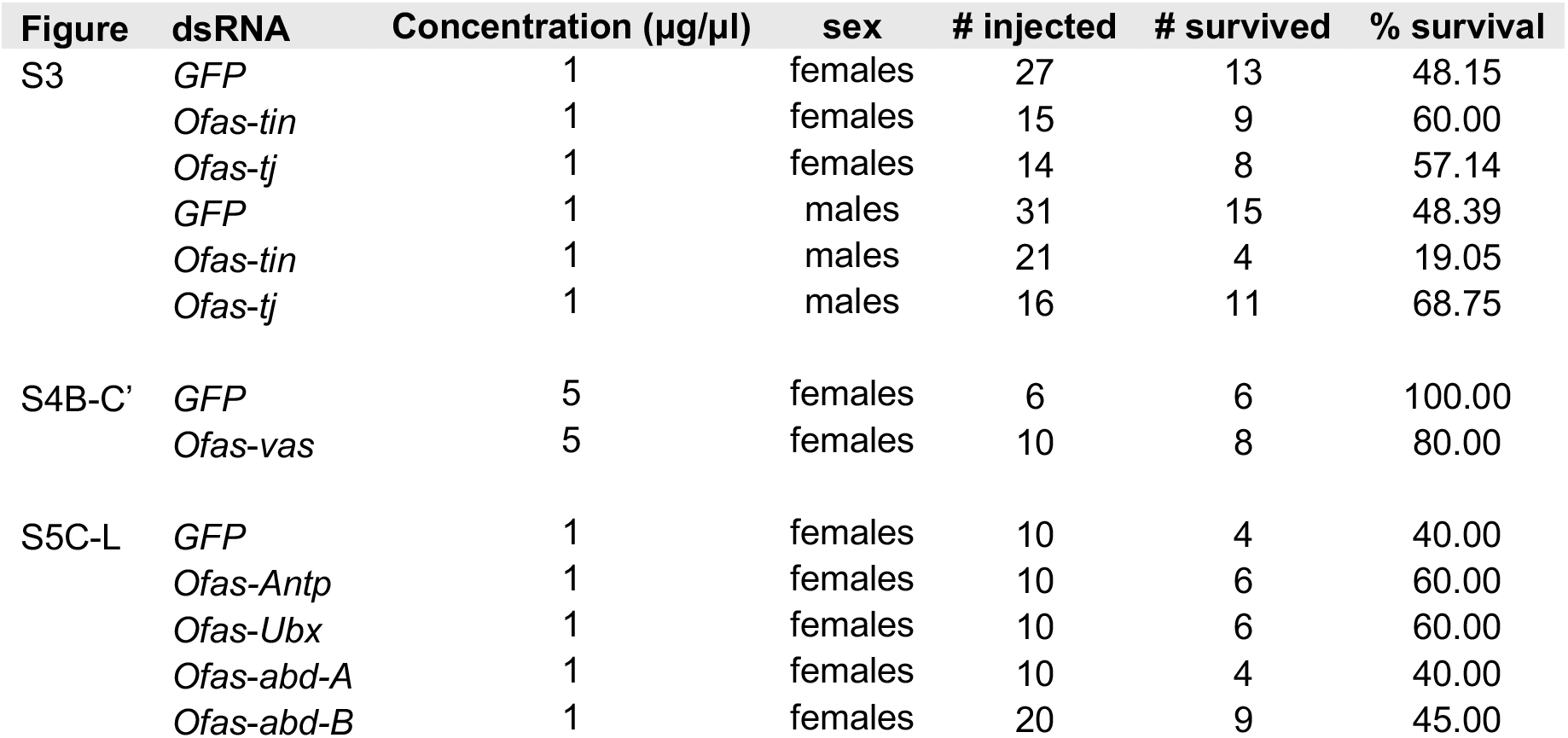
Adult injection statistics. Survival statistics for adult injection experiments. An individual was scored as “survived” if it was still living ten days after injection.

**Table S8.**
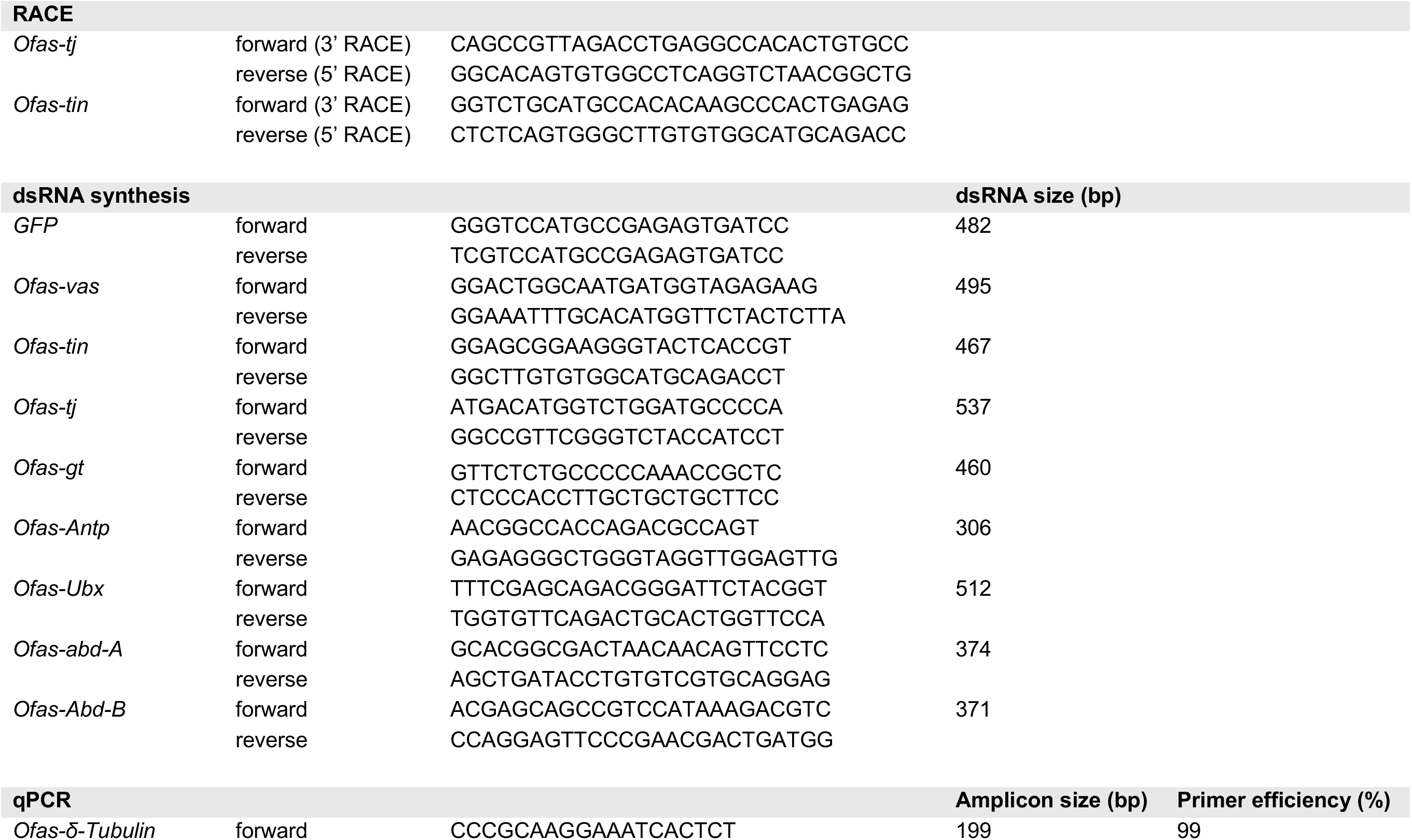
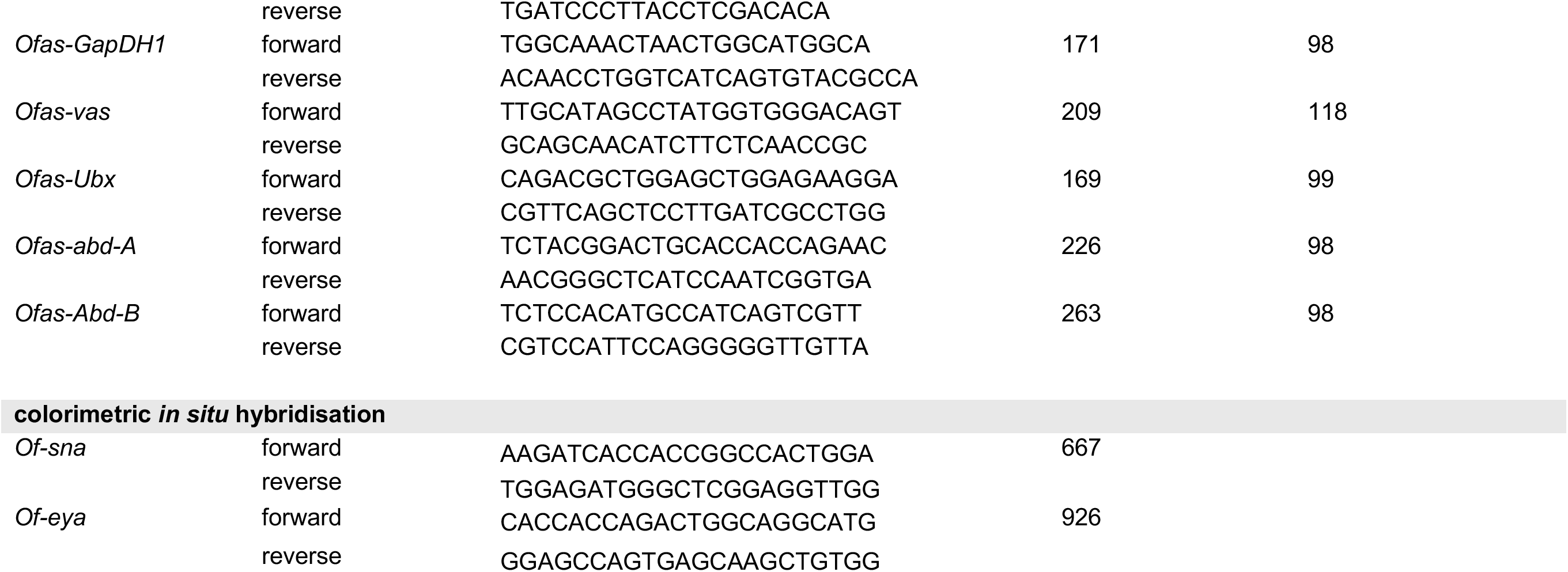
Primers used. Primers used in this study for experiments indicated in the headers. qPCR amplification efficiency for each primer pair was calculated from the slope of a standard curve generated by plotting C_g_ values against the logarithm of the input cDNA concentration for four serial dilutions ^211^.

**Table S9.**
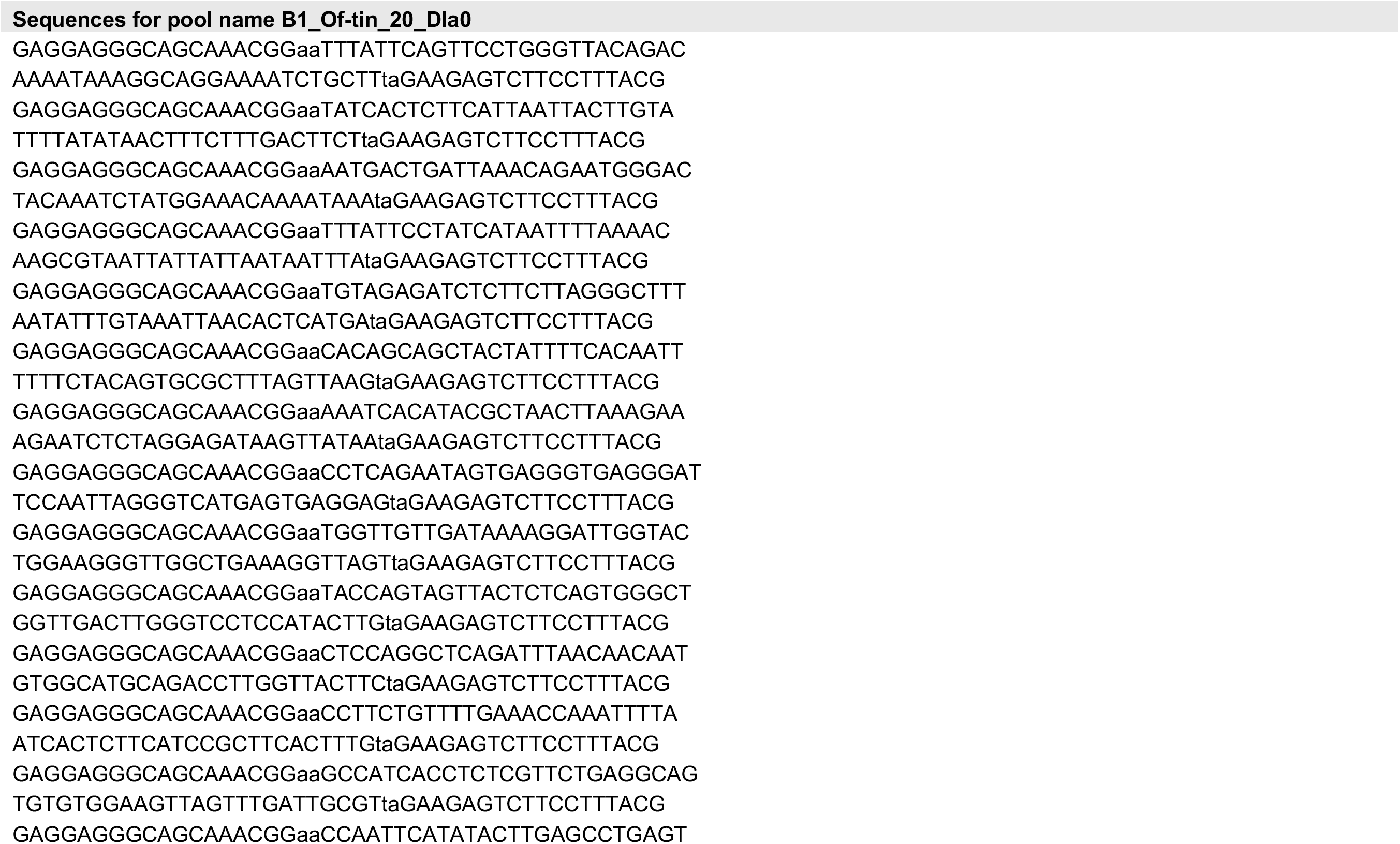
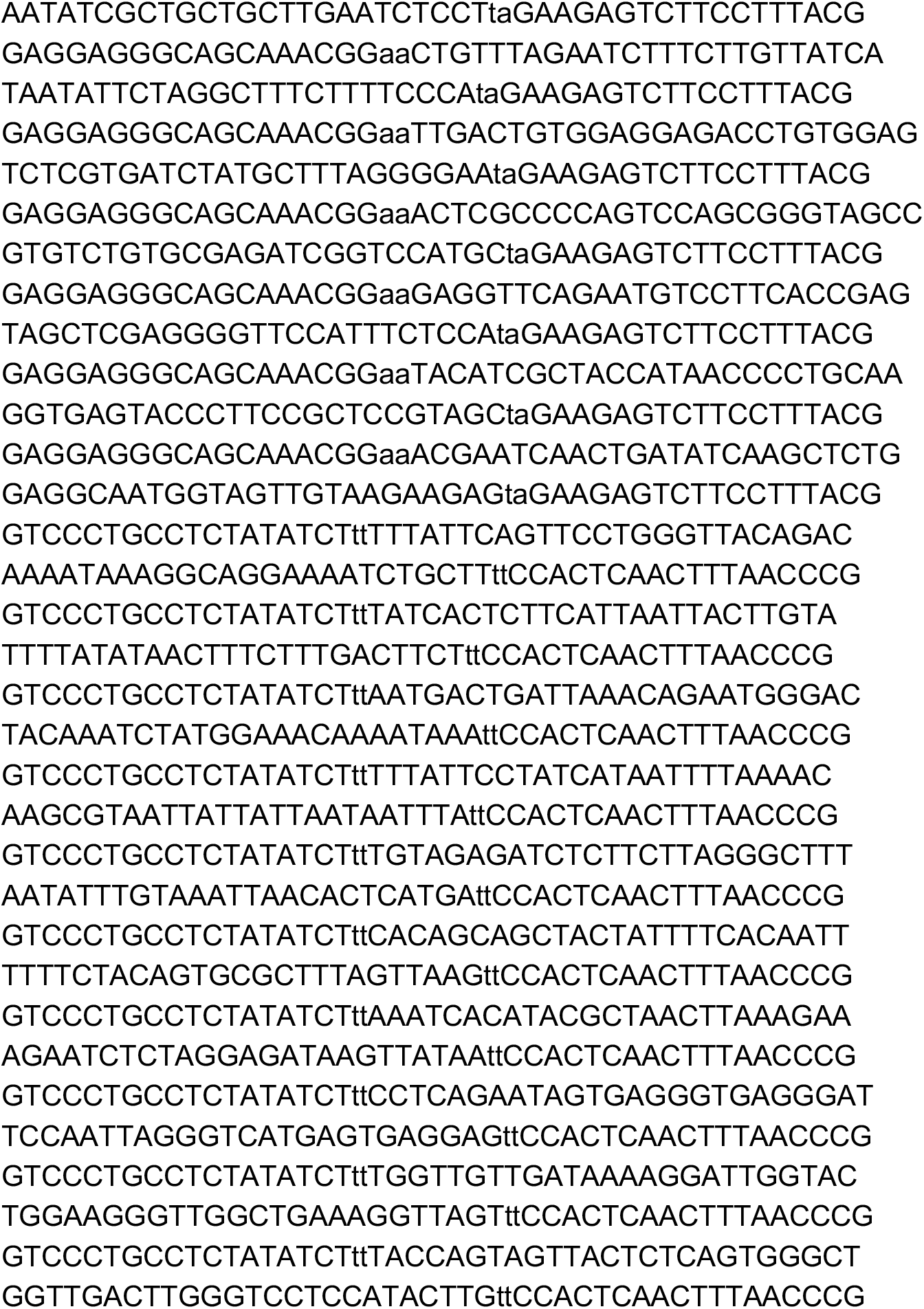
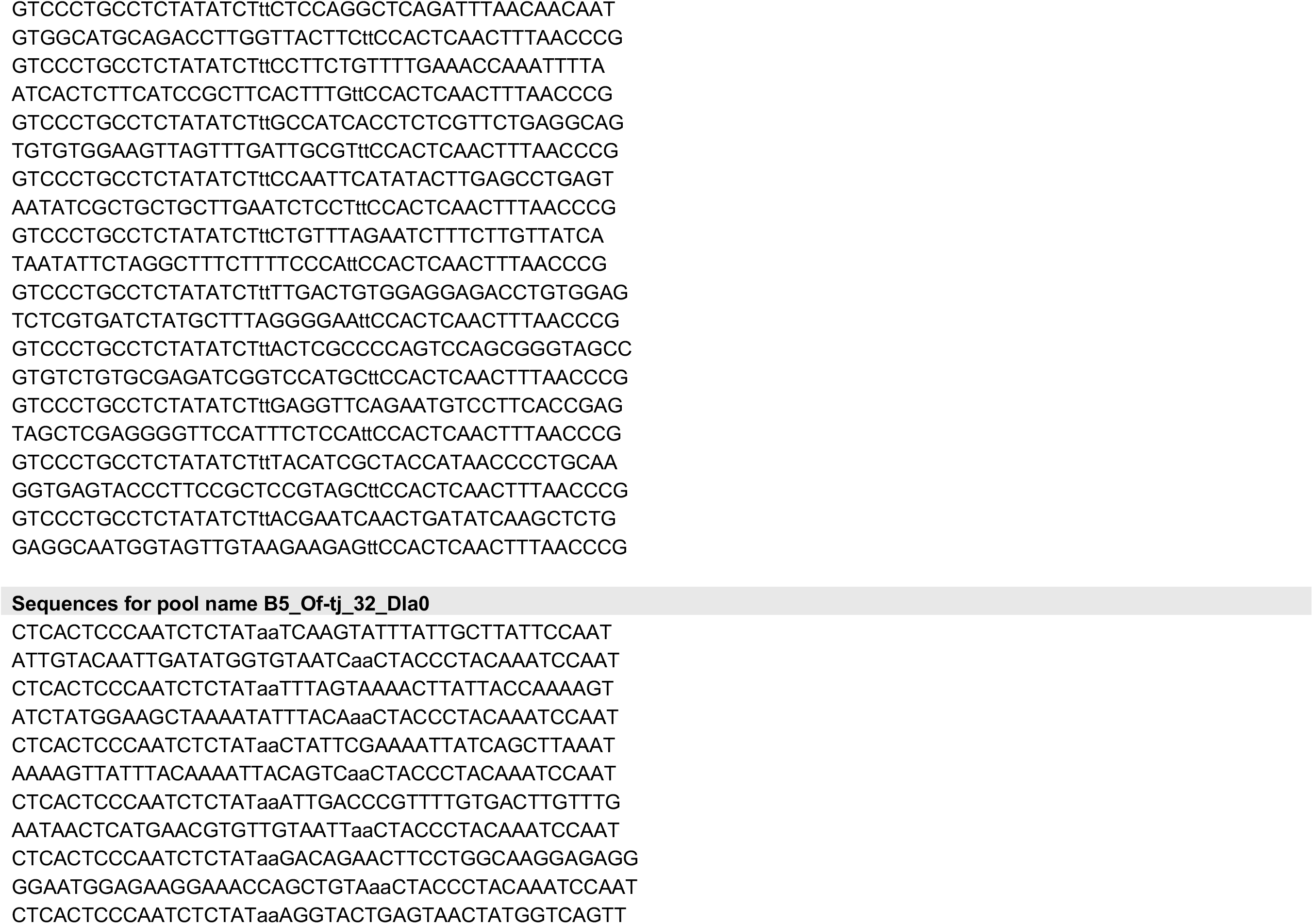
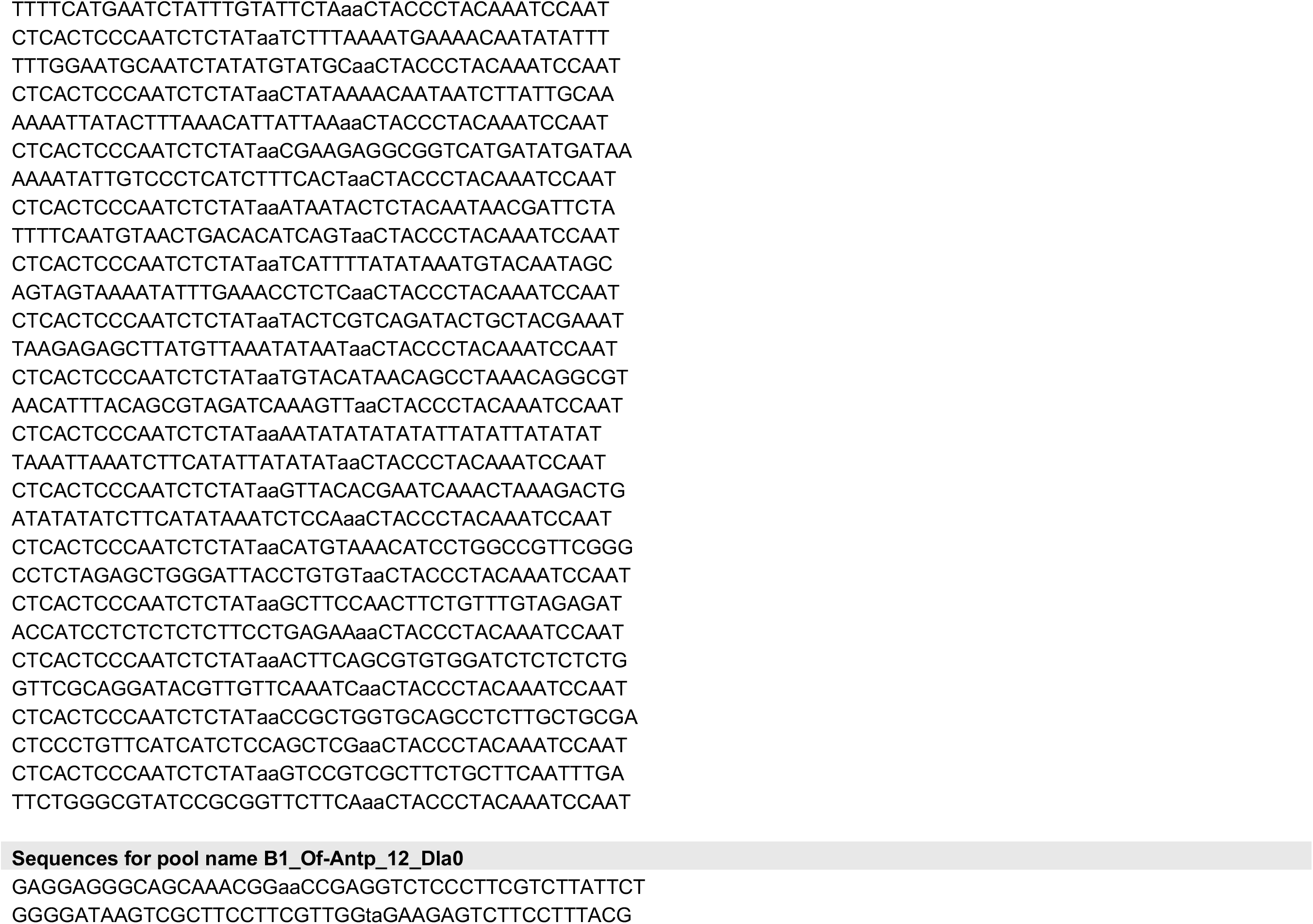
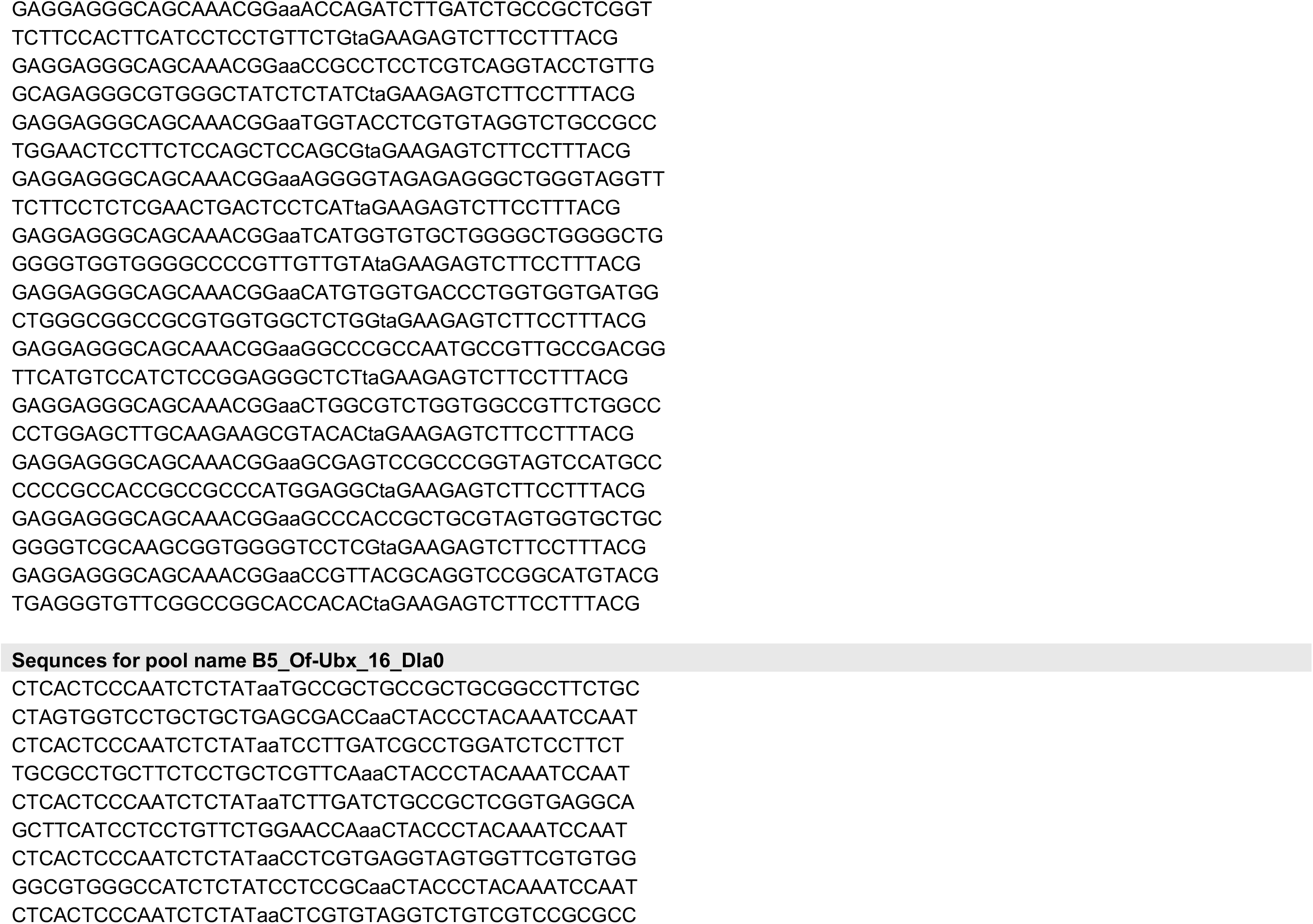
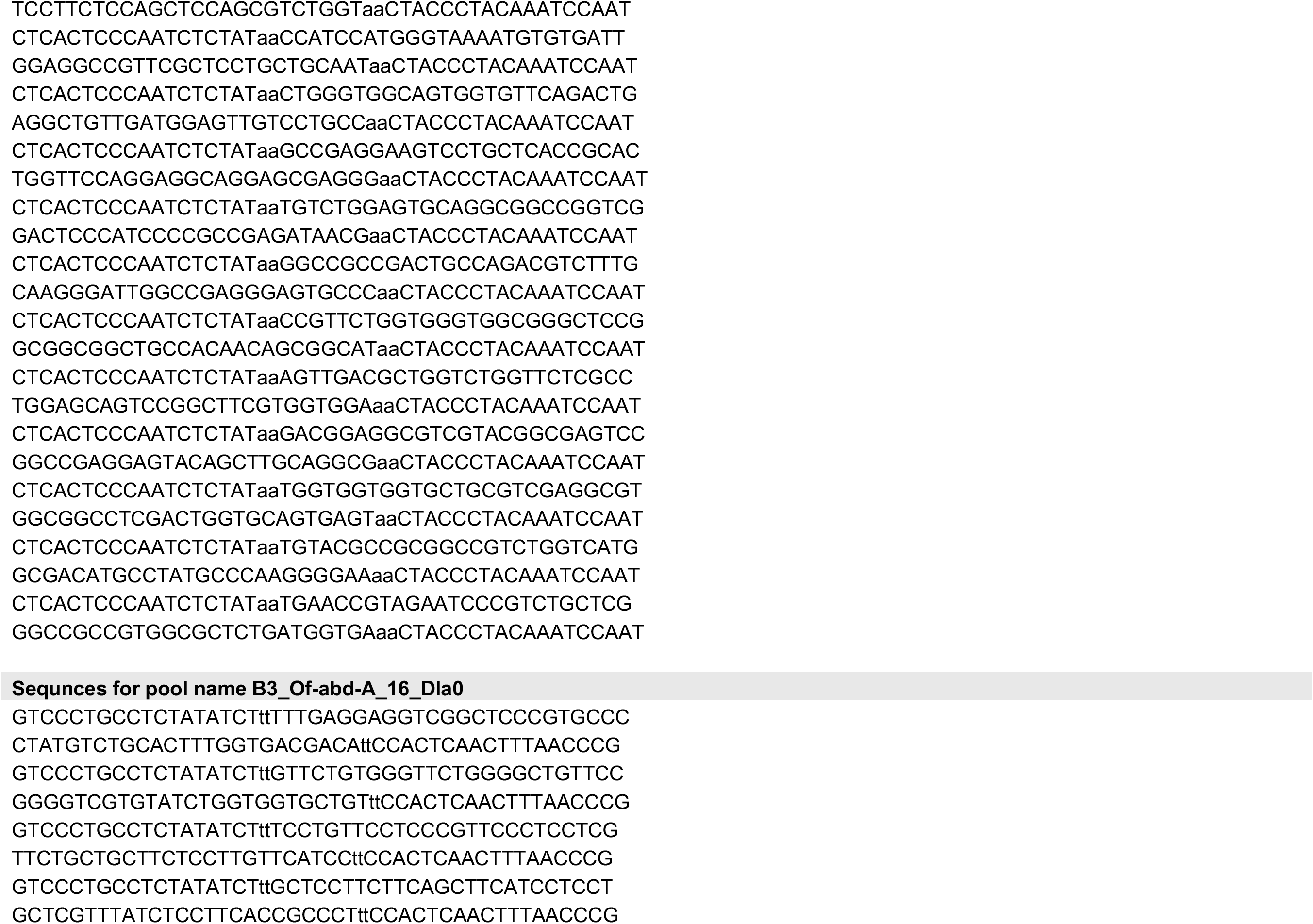
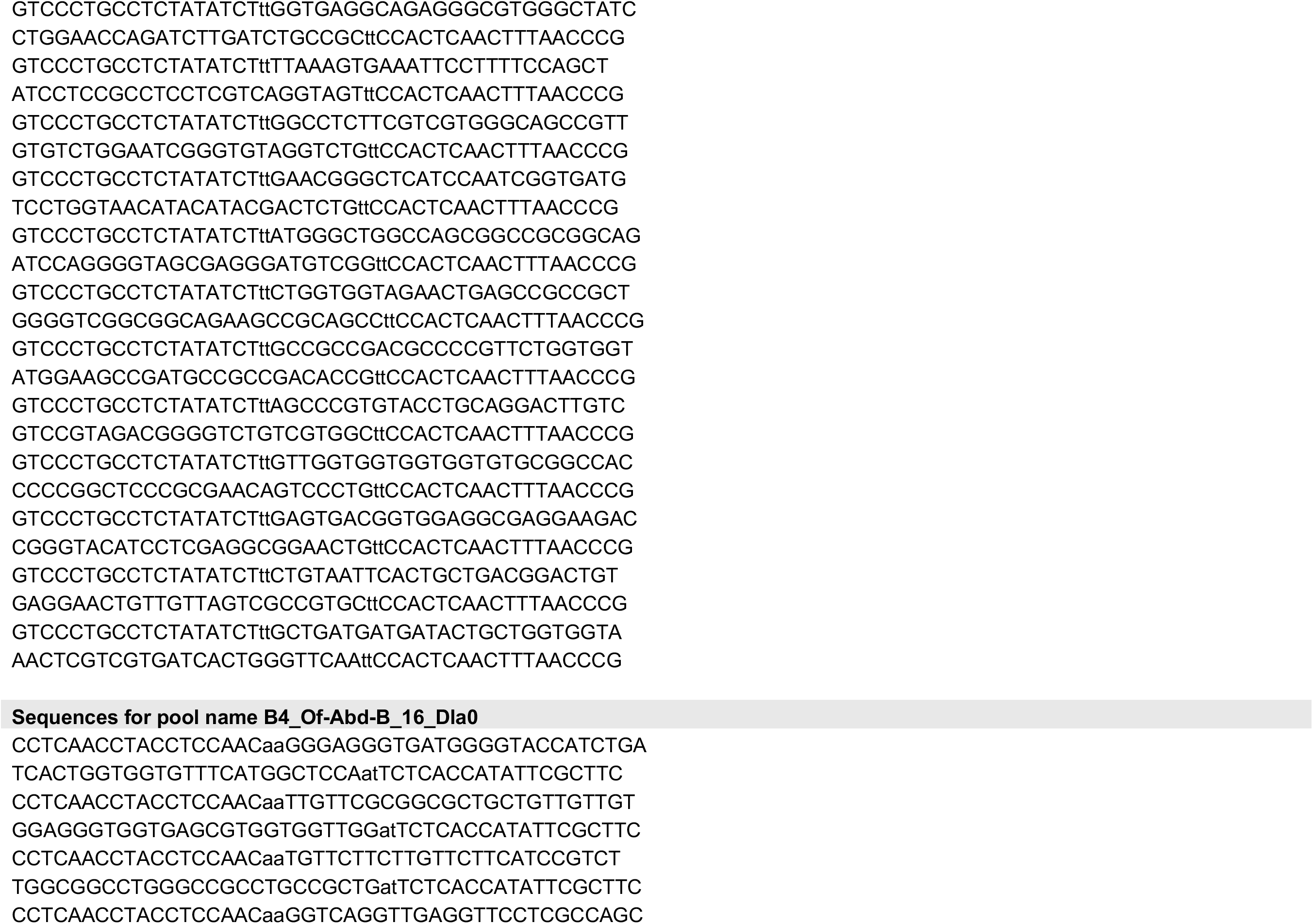
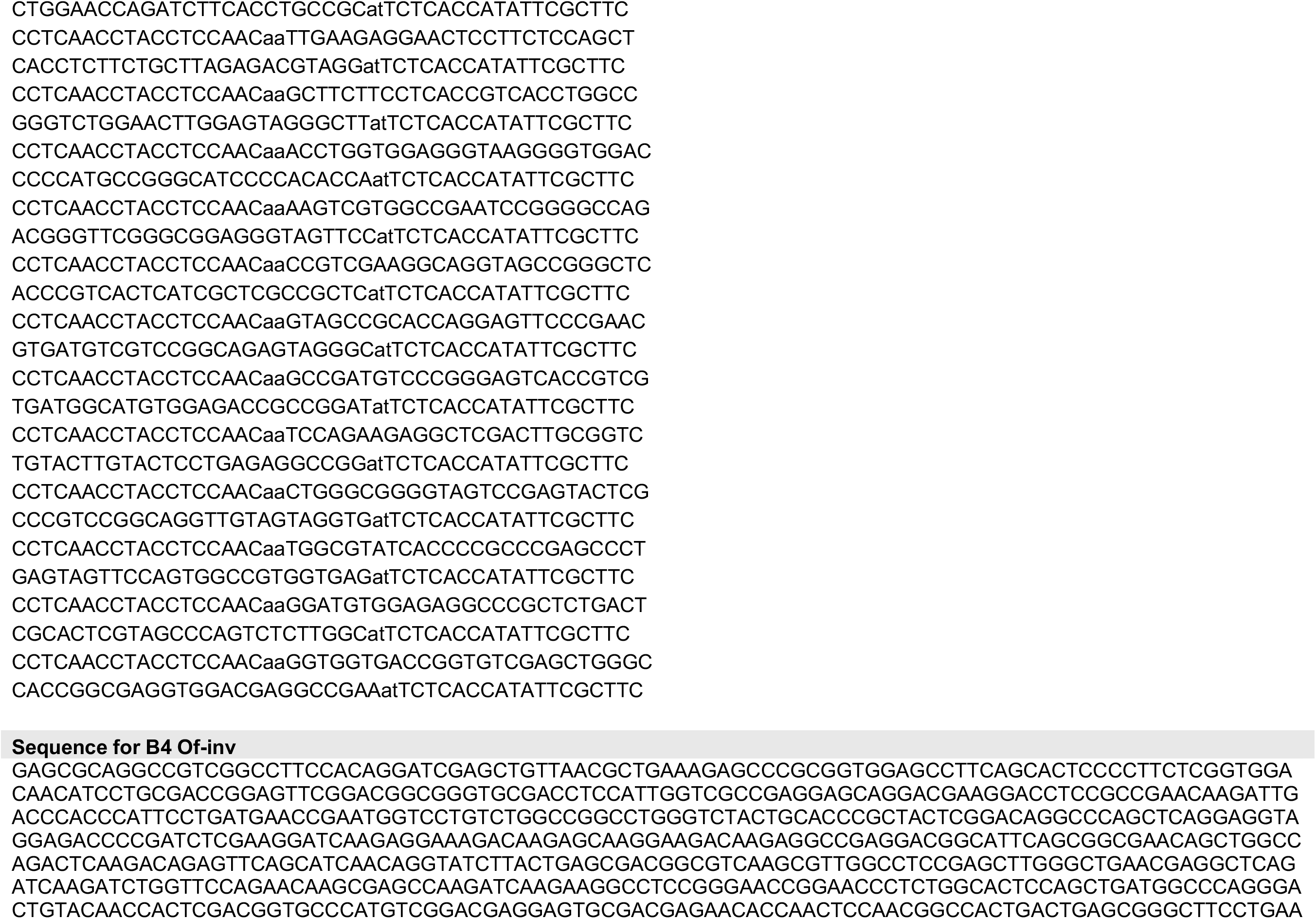
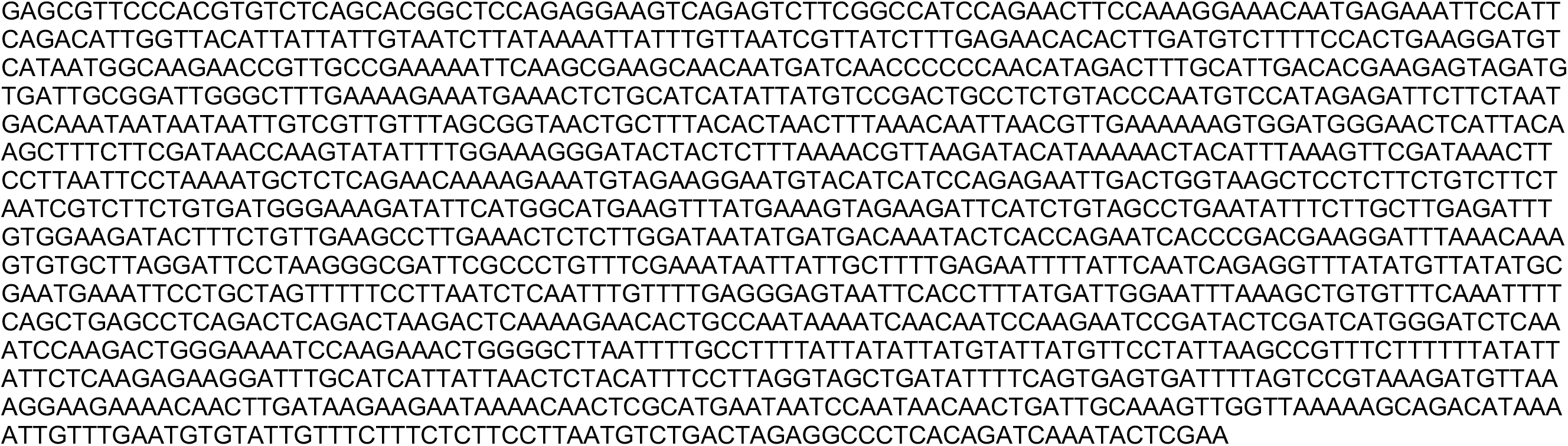
Sequences of oligos used in hybridisation chain reaction (HCR) experiments. All HCR probes except for that for *Ofas-inv* were designed using HCR Probe Maker (Özpolat lab, Washington University Saint Louis ^167^) and ordered from Integrated DNA Technologies as oPools (oligo pools). The *Ofas-inv* probes were obtained from Molecular Instruments by submitting a partial cDNA sequence (shown below).

